# Eliminating Genes for a Two Component System Increases PHB Productivity in *Cupriavidus basilensis* 4G11 Under PHB Suppressing, Non-Stress Conditions

**DOI:** 10.1101/2022.09.19.508597

**Authors:** Kyle Sander, Anthony J. Abel, Skyler Friedline, William Sharpless, Jeffrey Skerker, Adam Deutschbauer, Douglas S. Clark, Adam P. Arkin

**Affiliations:** Center for the Utilization of Biological Engineering in Space, Berkeley, CA; Department of Bioengineering, University of California, Berkeley. Berkeley, CA; Department of Chemical & Biomolecular Engineering, University of California, Berkeley. Berkeley, CA; Environmental Genomics and Systems Biology Division, Lawrence Berkeley National Laboratory, Berkeley, CA; Department of Bioengineering, University of San Diego, San Diego, CA; Genome Science and Technology Graduate Program, University of British Columbia, Vancouver, BC, Canada; Zymergen, Emeryville, CA

**Keywords:** PHB, Bioplastic, Cupriavidus, RB-TnSeq, Histidine Kinase

## Abstract

Species of bacteria from the genus *Cupriavidus* are known, in part, for their ability to produce high amounts of poly-hydroxybutyrate (PHB) making them attractive candidate bioplastic producers. The native production of PHB occurs during periods of metabolic stress, and the process regulating the initiation of PHB accumulation in these organisms is not fully understood. Screening an RB-TnSeq transposon library of *Cupriavidus basilensis* 4G11 allowed us to identify two genes of an apparent, uncharacterized two component system which, when omitted from the genome, are capable of increased PHB productivity in balanced, non-stress growth conditions. We observe average increases in PHB productivity of 56% and 41% relative to the wildtype parent strain, upon deleting each of two genes individually from the genome. The increased PHB phenotype disappears, however, in nitrogen-free unbalanced growth conditions suggesting the phenotype is specific to fast-growing, replete, non-stress growth. Bioproduction modeling suggests this phenotype could be due to a decreased reliance on metabolic stress induced by nitrogen limitation to initiate PHB production in the mutant strains. Such strains may allow for the use of single stage, continuous bioreactor systems, which are far simpler than PHB bioproduction schemes used previously. Bioproductivity modeling suggests that omitting this regulation in the cells may increase PHB productivity up to 24% relative to the wildtype organism when using single stage continuous systems. This work furthermore expands our understanding of the regulation of PHB accumulation in *Cupriavidus*, in particular the initiation of this process upon transition into unbalanced growth regimes.

## Introduction

Bioderived plastics offer a potentially sustainable and renewable alternative to conventional petroleum-derived plastics. Widespread production and use of bioproduced plastics remain limited, in part, by low productivies^1–3^. Species of the bacterial genus *Cupriavidus* exhibit the highest productivities of natural or recombinant poly-hydroxybutyrate (PHB) producing organisms^4^. While it is possible to attain very high PHB productivities using a variety of organisms through sophisticated multistage bioreactor or feeding regimes^5–7^, these systems can require very high substrate loadings, suffer from low overall substrate utilization, and necessitate regular shutdown and start-up cycles in-between batches^4,5,8,9^. Maintaining high substrate utilization remains important for both economically viable industrial bioplastic production^2^ and in resource-constrained, remote and austere bioplastic production and deployment settings^10^.

PHB is natively produced in *Cupriavidus* during conditions when the organism is incurring any of a variety of metabolic stresses and the productivity potential in *Cupriavidus* is typically maximized by initiating high PHB productivity via one or several of these stresses after a period of cell biomass production. This strategy typically requires multi-stage sophisticated bioprocesses^11,12^. Though relatively high PHB productivities have been achieved, intracellular PHB accumulation during non-stressed, balanced growth remains low (~<50% PHB cdw basis) and thus there remains further potential to increase PHB accumulation in the organism under non-stress conditions, fast-growing, replete conditions.

We propose overall productivity might be increased further by engineering *Cupriavidus* to produce higher amounts of PHB continuously, during periods of balanced growth. Organisms which produce PHB using recombinant genes to produce PHB do so in an apparently constitutive manner^13^. One of the highest productivities achieved in *Cupriavidus* was achieved by allowing a slight amount of residual growth, while still maintaining nutrient limited conditions^14^, suggesting that growth-associated PHB production is not only possible but is a promising way to rapidly increase overall PHB productivity. Similarly, elemental mode analysis suggests the possibility and promise of constitutive PHB bioproduction by *Cupriavidus*^15^.

Only segments of the genetic and metabolic mechanisms regulating PHB production in *Cupriavidus* have been elucidated, and engineering genes of the PHB production pathway (*phaCAB*) directly has yet to enable growth-associated PHB production in *Cupriavidus*. This suggests its productivity is regulated by availability of redox cofactors^14^ and/or determined by regulatory elements^16–18^.

Enforcing growth-associated bioproduction has been a strategy for increasing productivity for a number of products and host/chassis organisms^19^ and for PHB productivity in *Synechosystis*^20^. This generic strategy considers only augmentation of metabolic genes and often requires tenuous and unreliable manipulation of many genes in a single strain. This analysis can also indicate that growth-coupled production of a particular product is simply not possible given the metabolic systems considered. Alternatively, eliminating native regulatory genes from the genome has been shown to increase PHB productivity in *Cupriavidus* during conditions of minimized oxygen stress^21^.

In this work, we aim to genetically engineer PHB producing *Cupriavidus basilensis* 4G11, for constitutive bioproduction of PHB. The multiply-regulated nature of PHB synthesis^16–18,21^ suggests a whole-genome wide search for engineering targets is appropriate for gene targets regulating PHB productivity or associated gene-expression, whose absence could enable growth-associated PHB productivity. For this reason, we chose to screen an existing RB-TnSeq transposon library of *Cupriavidus basilensis* 4G11^22^ for mutants exhibiting constitutive bioproduction of PHB. *C. basilensis* 4G11 is closely related to the well-studied and industrially relevant PHB producing strain *C. necator* H16; with 79.5% of *C. basilensis* 4G11 proteins having one or more homologs in *C. necator* H16 (figure S1). *C. basilensis* 4G11 was originally isolated from subsurface groundwater at the Field Research Center at Oak Ridge National Laboratory which contains legacy contaminants originating from the production of nuclear materials^23^.

We adapted and used a FACS-based screen for intracellular PHB which utilizes BODIPY 493/503 dye^24^ (figure 1), and developed a culture pre-conditioning protocol which significantly suppressed/eliminated PHB productivity (figure S2). PHB productivity in candidate mutants was then measured directly in these same conditions and shown to be higher, suggesting a role for these genes in regulating the onset of PHB production in *Cupriavidus*, and showing that the elimination of these genes may enable constitutive PHB bioproduction.

**Figure 1:**
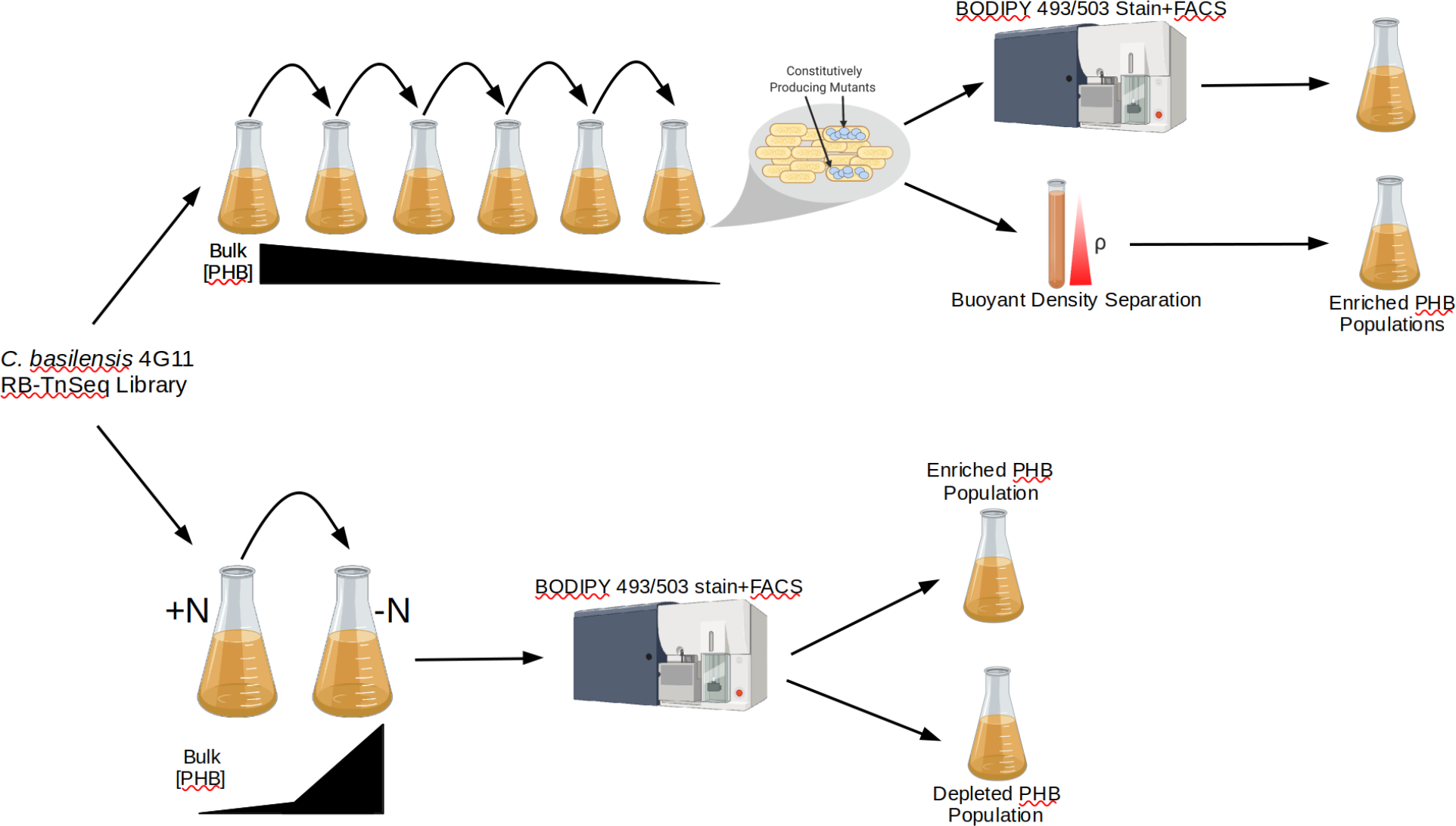
Dual screening strategy utilized in this study to isolate mutants exhibiting increased PHB accumulation phenotype from *C. basilensis* 4G11 RB-TnSeq libraries. Cells are repeatedly subcultured in replete medium to maximally suppress PHB accumulation among the bulk population, shown along top of the figure. From this population, high-PHB accumulating (‘constitutively producing’) mutants were selected using two different fractionation methods: FACS and buoyant density separation. Nitrogen-starved cultures were first grown in replete medium, followed by an incubation period in nitrogen-free medium in which the bulk population accumulates high amounts of PHB, shown along the bottom of the figure. From this population, we separately selected the highest and lowest accumulating PHB mutants.

## Results

### Development of Screen to Minimize/Suppress PHB Production in Wildtype Organism

In our screen, we sought mutants which accumulate differentially more PHB in non-stress, highly replete, fast growing conditions; conditions in which wildtype species of *Cupriavidus* and other PHB accumulating organisms accumulate very little, if any, PHB. To assess this specific phenotype, we developed a method for culturing *C. basilensis* 4G11 which reliably suppressed intracellular PHB accumulation to a minimal amount. This method resulted in cells in culture that were in a fast-growing, nutrient-replete state, as we see from the reproducible near-complete suppression of intracellular PHB (figure S2). This was achieved through frequent passaging of cultures, maintaining minimal culture OD, maximizing nutrient concentrations and maintaining saturated dissolved oxygen conditions. This method for suppressing intracellular PHB was used prior to both RB-TnSeq library screens as well as before and during quantification of PHB productivity in candidate mutants. In practice, this minimized potential background PHB, and served as a non-stressed, nutrient-replete state in which we performed PHB quantification in candidate mutants for constitutive PHB bioproduction.

### Screening RB-TnSeq Libraries for High-Producing Mutants

We performed forward genetic screens on a pre-existing *C. basilensis* 4G11 RB-TnSeq library^22^ for increased PHB productivity under both PHB-suppressed replete conditions (as described in the previous section), and in conditions inducing high PHB accumulation; through incubation in medium in which all nitrogen had been omitted after a period of initial cell growth in nitrogen-replete medium (figure 1). We identified eight candidate genes whose transposon insertion mutant strains showed increased enrichment when screened for increased intracellular PHB in either replete, fast-growing conditions or nitrogen-starved conditions (Table S1). Several genes identified were genomically co-localized, suggesting possible inter-operonic expression and/or related metabolic function; specifically RR42_RS17055 and RR42_RS17060 form two genes of an apparent two-component histidine kinase/response regulator system, while RR42_RS18960-RR42_RS18970 are genes involved in membrane asymmetry maintenance. The identified collocated genes also exhibited consistent gene fitness (figure S3) in several previously conducted stress exposure or defined nutrient auxotrophy assays^22^ further implicating their related and/or dependent gene function^25^.

### Assaying Candidate Mutants for Constitutive PHB Productivity

We generated markerless whole-gene deletion mutants for identified genes to quantify their PHB productivity. We utilized genetic and conjugation methods developed previously for *Cupriavidus necator*^18,26^. Similar to *Cupriavidus necator*, *C. basilensis* 4G11 was found to be gentamicin resistant (data not shown). For this reason, to make deletion mutants we utilized suicide vectors based on the pMQ30 backbone^27^ containing a kanamycin resistance marker in place of the gentamycin resistance marker on the original pMQ30 plasmid. We removed the heterologous kanamycin resistance using the *sacB* gene and sucrose counterselection. These mutants constitute, to our knowledge, the first markerless deletion mutants in this species.

We assayed these mutants for PHB productivity under fast-growing replete conditions. We found two mutants which consistently accumulated high amounts of PHB (figure 2) in this balanced-growth condition; a deletion mutant of the gene RR42_RS17060 which exhibited an average 56% higher PHB productivity relative to the wildtype parent strain, and a deletion mutant of the neighboring gene RR42_RS17055 which exhibited an average 41% higher PHB productivity relative to the wildtype parent strain. While mutants in other genes exhibited increased PHB productivity in single subcultures, no other genes assayed exhibited consistent productivity increase across subsequent subcultures as these two genes did.

**Figure 2:**
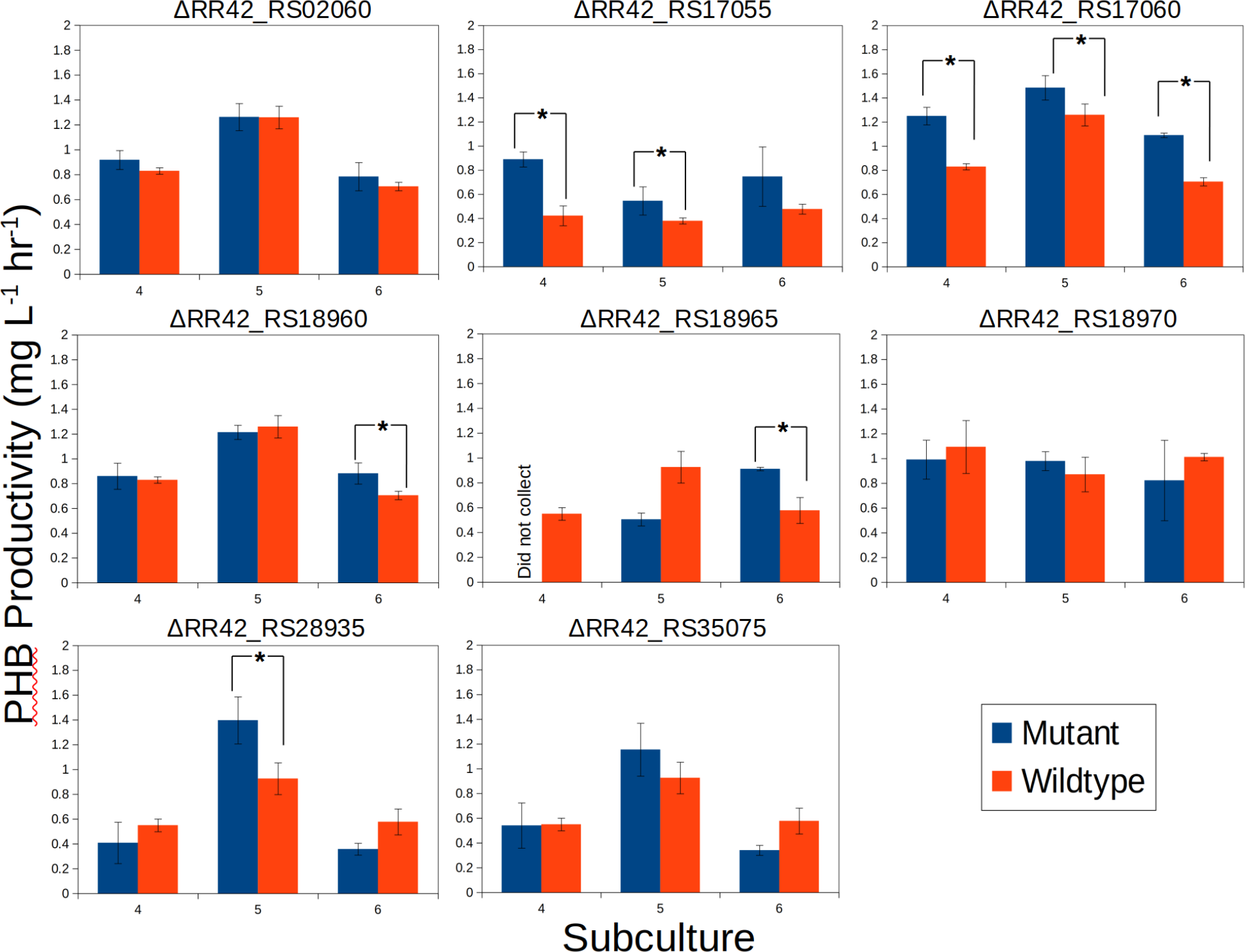
Accumulation of PHB in deletion mutants of selected genes during 4-6th subcultures in balanced-growth conditions. Genes encoding an apparent uncharacterized two-component system (RR42_RS17055-17060) consistently accumulate more PHB than the wildtype parent strain in non-stress conditions in which PHB accumulation is suppressed. * denotes statistical significance (one tailed t-test, n=3, equal variance, p<0.05).

This phenotype of mutants in these two genes appears to be specific to high-growth, non-stress, replete conditions. The PHB accumulation advantage disappears in ΔRR42_RS17055 and ΔRR42_RS17060 upon allowing replete, defined medium growth/incubations to continue to stationary phase, and when the same cells are subsequently incubated in defined minimal medium containing no added source of nitrogen (figure S4). The wildtype organism accumulates more PHB than either mutant upon reaching stationary phase after a 24 hour growth period in replete, defined medium, and more than the ΔRR42_RS17060 mutant after a subsequent 24 hour incubation in nitrogen free, defined medium.

RR42_RS17060 is annotated as a hybrid histidine kinase and RR42_RS17055, annotated as a LuxR family transcription factor, is likely the cognate response regulator for the histidine kinase as it contains a response regulator domain within its coding region (figure 3). Similar fitness patterns between these two genomically adjacent genes (figure S3) suggest they likely form a two-component signal sensing and transduction system (TCS). RR42_RS17055 also contains a DNA binding domain, suggesting it likely regulates transcription of genes it regulates in response to signals sensed by the histidine kinase. Gene annotations do not suggest any information about the external stimulus that the system responds to, or the genes it regulates the expression of. Promoter alignments of these two genes and the two additional genes which exhibit cofitness do not yield consensus sites indicative of potential operator sequences (data not shown).

**Figure 3:**
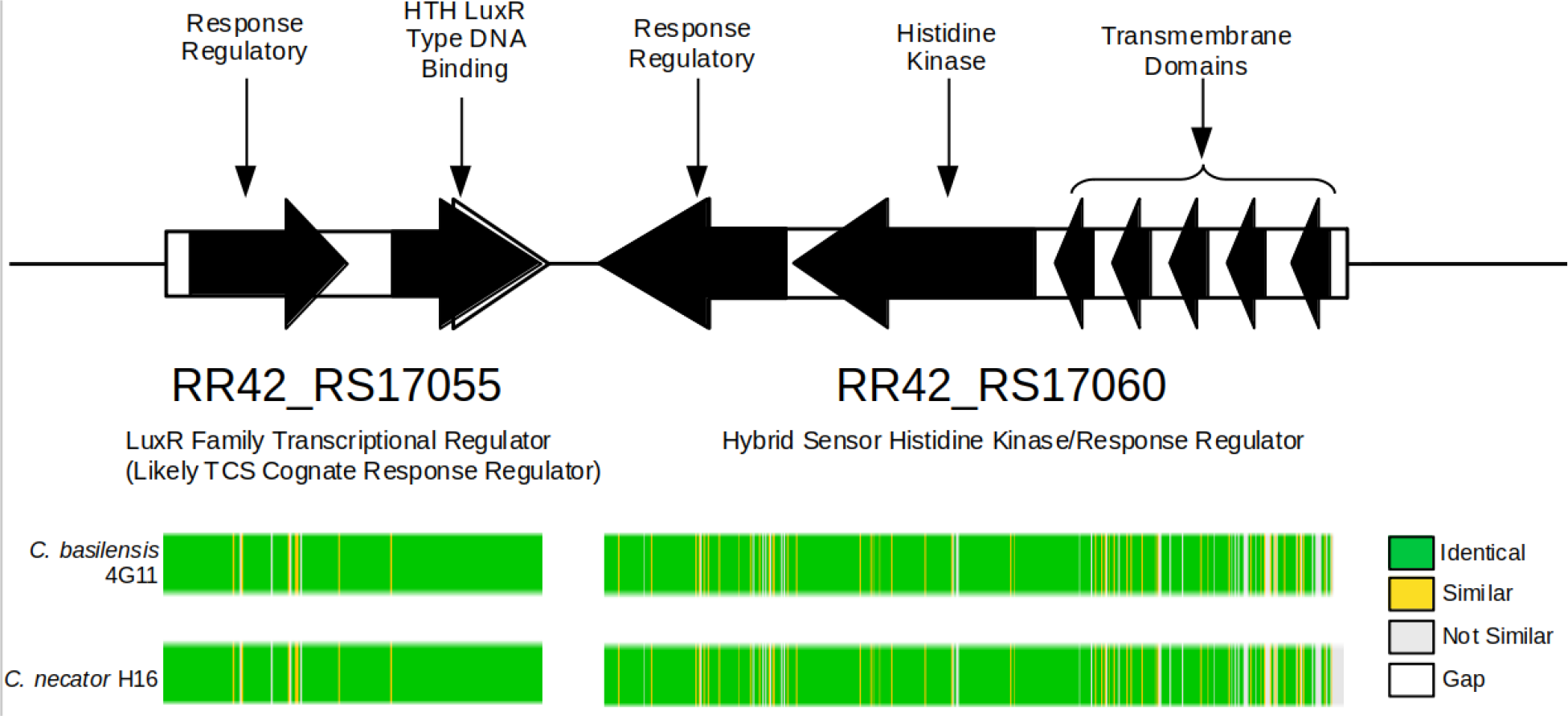
Genomic region, gene and domain orientation of genes comprising the two-component system in *C. basilensis* 4G11. RR42_RS17060 is annotated as a hybrid histidine kinase due to the presence of both histidine kinase domains and a response regulatory domain. RR42_RS17055 also contains a response regulatory domain as well as a DNA binding domain. Gene fitness similarities between the two genes (shown in figure S3) suggest they transduce the same signal(s) and the expressed proteins interact with each other. The presence of a DNA binding domain in RR42_RS17055 suggests signal transduction results in changes in DNA transcription. Colored bars indicate reciprocal amino acid homology between proteins encoded by these genes and their corresponding homologs encoded in the genome of *C. necator* H16. Whole protein homology between the two organisms is 95% and 84% for the LuxR Family Transcriptional Regulator and Hybrid Sensor Histidine Kinase/Response Regulator, respectively.

### Productivity Modeling Indicates TCS Mutants May Increase PHB Productivity

We built and interrogated a PHB kinetic model for two purposes: to confirm our hypothesis that this increase in PHB could be due to a decrease in reliance on unbalanced growth for the initiation of PHB production, and to decipher potential increases in PHB productivity that might be realized using these strains in a continuous bioproduction system.

We adapted a PHB production model developed previously^15^ which has been utilized to model PHB production dynamics in *C. basilensis* 4G11^28^. This kinetic model considers cell biomass growth, production of PHB, consumption and cell growth inhibition of key metabolites acetate and nitrogen (in the form of ammonium chloride). It models the coupling of nitrogen concentration and PHB productivity through the key model parameter K_PIN_, which represents the sensitivity and reliance of PHB accumulation to the nitrogen concentration for both wildtype and mutant strains. Our model fit for this parameter of the wildtype parent strain was nearly the same as predicted in a previous study utilizing the same growth medium through different culturing conditions^28^. We further recapitulated similar cell and PHB substrate yield values to known, previously calculated values (table S2).

In fitting the K_PIN_ parameter to our cell growth and PHB production data from single deletion mutants of the RR42_RS17055-17060 two component system, we found this parameter to be much higher in these mutant strains than it was in the wildtype (table S2). This larger value of the parameter might be interpreted as our mutants’ decreased reliance on nitrogen limitation metabolic stresses to initiate PHB accumulation. While our adaptation of this model assumes PHB accumulation is only coupled to nitrogen concentration, it is known that many metabolic stresses and nutrient deficiencies can initiate PHB production in the closely related *Cupriavidus necator*^15^, and this may also be the case for *C. basilensis* 4G11 as well. The higher value of K_PIN_ suggests that the two single deletion mutants (RR42_RS17055, RR42_RS17060) have PHB production metabolisms that are less coupled to nitrogen limitation or, perhaps more generally, to metabolic stress or nutrient deficiency, though this remains to be further explored or demonstrated.

We next developed a continuous bioproduction model to ascertain relevant and realistic bioproduction increases that might be realized by our single strain deletion mutants. We modeled PHB productivity of the wildtype strain and the ΔRR42_RS17060 using key model parameters carried forward from the previous analysis. We modeled PHB production in a single-stage continuous system as this system has the distinct advantage of being simpler in configuration and operation than many of the more sophisticated multi-stage systems required to achieve high PHB productivity in native non growth-associated wildtype *Cupriavidus necator*^6,11^. In our simulated continuous system the mutant is capable of 24% higher PHB productivity, generating a maximum of 0.541 g/L/hr in optimized conditions (figure 4a), compared to 0.437 g/L/hr produced by the wildtype *C. basilensis* 4G11 (figure 4b) in optimized conditions. Maximum mutant PHB productivity also exceeds wildtype productivities modeled over a range of dilution rates for both single and two-stage continuous systems (figure S5). Also, we observe an increase in the range of dilution rates and feed ammonia concentrations that would allow for near-optimal PHB productivity, relative to the wildtype organism.

**Figure 4:**
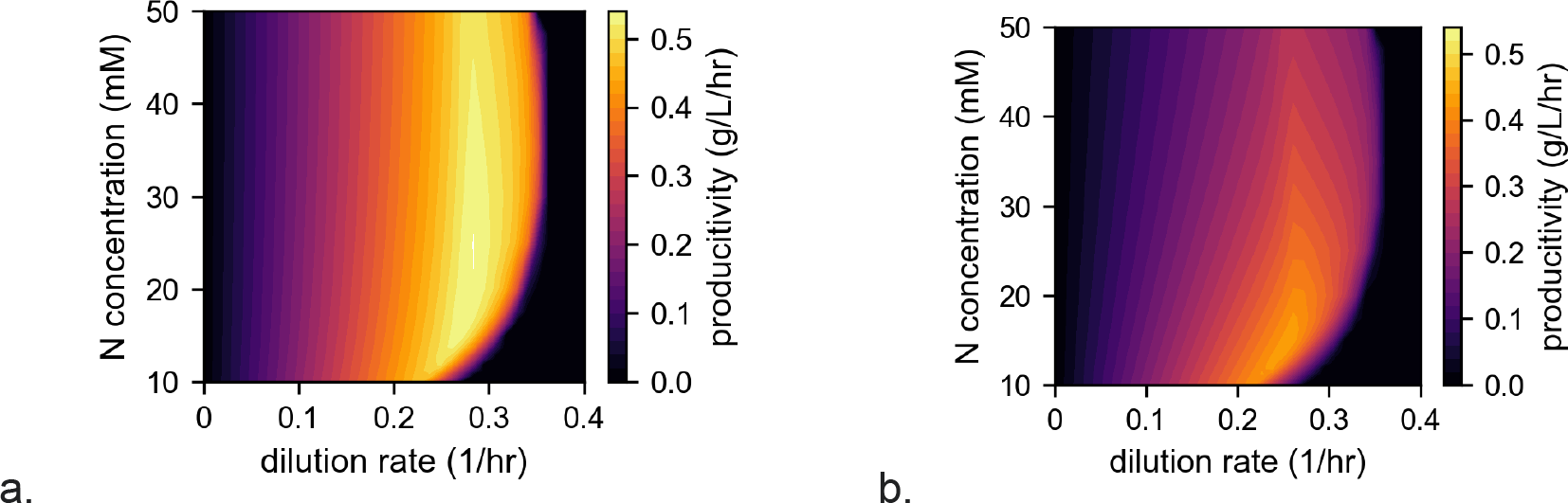
Volumetric PHB productivity of a) ΔRR42_RS17060 mutant and b) wildtype *C. basilensis* 4G11 in a simulated single-stage continuous system.

## Discussion

In this work we use a forward genetic screen alongside a novel replete-condition PHB productivity assay to identify two genes whose omission from the genome results in increased PHB productivity under non-stress conditions. These increases in productivity may translate to benefits in larger bioproduction settings given the possible use of simpler reactor configurations and operating needs relative to other multi-stage, batch configurations investigated previously. Also, an apparent decrease in necessary reliance on metabolic stress may simplify operational burden of far simpler reactor systems such as single stage continuous systems.

There is typically an inherent tradeoff between overall productivity and substrate utilization in producing microbial PHB^29^, and it is possible to achieve high PHB productivities at the expense of substrate utilization^30–32^. However there remain application scenarios where a high substrate utilization and yield must be simultaneously maintained; in particular when substrate costs are high and in remote and austere environments such as long term missions in space where access to feedstock material might be limited. In this work we fit an empirical substrate mass yield of 0.23 g PHB/g acetate (table S2), which coincidentally approximates those of commercial systems which predominantly use hexoses as substrates^33^. Though gene mutations identified in this paper may offer productivity increases to commercial systems without sacrificing substrate utilization, it remains unclear whether further improvements (through further metabolic engineering, optimizing cultivation conditions, etc.) will be needed for other production settings.

Methods for continuous, constitutive production of PHB in *Cupriavidus* have been proposed and sought previously^11^. In one instance PhaB was engineered to be NADH-dependent to couple with a recombinant NADH dependent glucose-6-phosphate to provide NADH demand^34^ and allow for fermentative, anaerobic PHB production without the need to inhibit biomass formation. This strategy was however not fully demonstrated, and may still be subject to gene regulatory constraints not considered. Many other metabolic systems previously shown to impact PHB accumulation in species of *Cupriavidus* were regulatory systems. This includes phosphoregulation^17,35^, the PhaR protein that binds upstream of and transcriptionally regulates the *phaCAB* operon^16,36^, PHB granule associated phasins^21,36,37^, and the stringent response^18,38^. The accumulation of PHB in *Cupriavidus* thus appears to be regulated in multiple ways and a comprehensive understanding of its regulation remains unknown, including any regulators which may sit atop known regulatory machinery in a regulation hierarchy. In this work we find another regulatory element capable of impacting PHB productivity, though the full scope of its regulatory sensing and action remains unknown. Knowing the regulon and metabolite(s) it responds to may allow for a more comprehensive understanding of the regulation of PHB synthesis in this and other organisms. Such knowledge may also suggest more effective methods with which to engineer increased PHB productivity, through deregulating the onset of PHB synthesis, in this and related organisms.

The two genes identified in this screen demonstrated to increase balanced-growth productivity are not functionally characterized. They likely comprise a two-component system responsible for sensing a metabolic signal and transducing a response signal through altering gene transcription. RR42_RS17060 is annotated as hybrid histidine kinase and contains domains associated with both histidine kinase activity and a response receiver/regulator in the same CDS (figure 3). The adjacent gene RR42_RS17055 also contains a response regulator domain, as well as an HTH DNA binding motif. A similar phenotype of knockout mutants of these genes suggests they are part of the same signal cascade. Two other genes share similar fitness phenotypes with these genes (figure S3) in an array of other assays performed previously^22^, though annotations of these genes do not readily indicate the cellular function this apparent system may be directly regulating, nor how this regulation may be otherwise impacting PHB accumulation in replete, high growth rate conditions. Additionally, the increased PHB accumulation phenotype exhibited by these two mutants is apparently specific to fast-growing, non-stress conditions. The disappearance of the increased PHB accumulation phenotype (figure S4) in more limiting and stress conditions suggests that the regulatory differences invoked by the two component system may be specific to the early onset of PHB accumulation in the organism and may be yet another part of the complex regulation of PHB accumulation in this and other species of *Cupriavidus*.

Cofitness analysis, made possible by another study using this same RB-TnSeq library^22^, indicates two other genes which show consistent fitness patterns with genes comprising this apparent two component system (figure S3). One gene is annotated as a Serine/Threonine Protein Phosphatase; RR42_RS04895. While the specific function or signal of the protein encoded by this gene remains unknown, this class of proteins is known to participate in phosphoregulation, through protein dephosphorylation activity, of stress-related signal cascades^39^. Similarly, the interaction partners or specific function of RR42_RS02045, annotated as a GTPase remain unknown. Notably, protein sequence homologs exhibiting >84% homology to all four proteins (the two collocated apparent TCS genes, Ser/Thr Protein Phosphatase, and GTPase) also exist in the genome of the well studied closely related notable PHB producer *C. necator* H16, suggesting these genes may also play a role in regulating the onset of PHB production in this organism as well.

Previously it was demonstrated that CrbR (part of the CrbR/CrbS two-component system) regulates expression of genes involved in acetyl-CoA metabolism in *Pseudomonas fluorescens*^40^, and the LuxR family transcription factor encoded by RR42_RS17055 is 34% homologous to CrbR in *P. fluorescens*. We find putative operator sites upstream of the *acs* (Acetyl-CoA Synthetase) gene, a gene in the CrbR regulon of *P. fluorescens*, across diverse species of *Cupriavidus* (figure S6). However, the same sites cannot be identified upstream of either RR42_RS17055 or RR42_RS17060. Furthermore, there is low whole-protein sequence homology between RR42_RS17060 and CrbS of *P. fluorescens*, further suggesting the two systems are not homologous.

RR42_RS17055-RS17060 also exhibit protein homology to VsrB and VsrC (60.1% and 62.1% identity, respectively) of *Ralstonia solanacaerum* ATCC 11696 across > 97% of the length of the protein coding sequences. These proteins also share strong domain and positional homology (data not shown) to the TCS genes identified in this study. VsrBC has been shown to regulate EPS I exopolysaccharide biosynthesis through regulating expression of genes of the EPS I gene cluster in *Ralstonia solanacaerum* AW^41–43^ and EPS I-derived exopolysaccharides have been shown to be the primary mechanism enabling plant virulence activity in the organism. It is not known whether *C. basilensis* 4G11 can colonize plant tissue, though other closely related species of *Cupriavidus* are known to do so^44,45^. VsrC binds a particular operator site with the consensus sequence GCGGGGGAA upstream of the EPS I biosynthesis gene cluster in *R. solanacaerum* AW and, in conjunction with XpsR, activates expression of the EPS I locus^41,43^. We were unable to identify this operator site in the homologous upstream region of the EPS I gene cluster in *C. basilensis* 4G11, or in other closely related species of *Cupriavidus*. Despite this, RR42_RS17055-RS17060 may be the VsrBC two-component system, perhaps binding at a different operator site sequence in *C. basilensis* 4G11 than that identified in *R. solanacaerum* AW, though we were unable to identify any candidate sites. VsrBC is not known to directly regulate PHB accumulation in *Cupriavidus* or any other species of bacteria, to our knowledge. If these genes are responsible for activating EPS I, eliminating either of these genes may facilitate such a carbon metabolism flux redirection toward PHB biosynthesis, yielding the phenotypes we observe. Alternatively, RR42_RS17055-RS17060 may regulate both EPS I expression/biosynthesis and PHB accumulation simultaneously.

To date, no engineering or interrogation of two component systems has been done in species of *Cupriavidus* for the purposes of increasing PHB or deciphering any TCS-related aspects of the native regulation of PHB synthesis/mobilization. However, overexpression of the native *E. coli* AtoSC two component system activating fatty acid biosynthesis genes in an *E. coli* mutant expressing *C. necator phaCAB* genes resulted in increased overall PHB titer and per-cell PHB accumulation^30^, linking native fatty acid metabolism to heterologous PHB production in the organism. More generally, TCS system engineering has resulted in bioproduction benefits for other products in other organisms as well such as the elimination of *gacS*-*gacA* system in *P. putida* resulting in an increased conversion of para-coumarate to indigoidine^46^. The successful leveraging of two component system engineering for bioproduction increase, and the identification of a two component system shown to impact PHB productivity in this study suggests these systems may be leveraged further in *Cupriavidus* to increase PHB productivity in both native and recombinantly producing organisms and to further elucidate the complex regulation of PHB productivity in these organisms^47^. While further molecular regulatory and mechanistic information about this two component system remains to be elucidated, utilizing these mutations to engineer bioproducing strains may allow for simpler, single stage continuous bioreactor settings to produce PHA using bacterial hosts. Similarly, engineering this system may relieve the constraint of needing to invoke nitrogen limitation or other stress, and the reliance on complicated, multistage bioreactor systems necessary for this, allowing instead the use of simpler continuous systems to achieve similar or greater PHA productivities.

## Conclusion

We identify genes encoding a two component system (RR42_RS17060; a hybrid histidine kinase and RR42_RS17055; a LuxR family transcription factor with a response regulator domain) capable of impacting PHB accumulation in *C. basilensis* 4G11. When each gene is individually omitted from the genome, these mutants are capable of producing higher amounts of PHB in balanced growth conditions (56% and 41% higher, respectively for ΔRR42_RS17060 and ΔRR42_RS17055). The absence of this phenotype in unbalanced growth conditions suggests these genes may impart their impact early in the process of initiating PHB accumulation in the cells and are yet another regulatory element shown to have an impact on PHB accumulation in species of *Cupriavidus*. Bioproduction modeling suggests mutants of these genes may be less sensitive to dependence on nitrogen limitation and possibly other metabolic stress than the wildtype organism to initiate PHB accumulation. Simulating PHB accumulation in an optimized simple, single stage continuous bioproduction setting suggests these mutants may enable PHB production increases of 24% over the wildtype parent strain.

## Methods

### Strains, Medium, Growth Conditions

*Cupriavidus basilensis* 4G11^23^ and previously-generated RB-TnSeq library generated for this organism^22^ were both maintained in R2A complex medium for general culture maintenance and storage. Experiments were performed in defined DM9 medium^48^ containing 2 g/L sodium acetate as the sole source of carbon and 1 g/L NH_4_Cl as the sole source of nitrogen, unless otherwise noted. Cultures containing the *C. basilensis* 4G11 RB-TnSeq library^22^ were grown in medium containing 100 ug/mL kanamycin sulfate (Millipore Sigma part number 60615-5G). Cultures were grown and incubated at 30°C shaking at 250 rpm in Ultrayield flasks. Cultures containing *C. basilensis* 4G11 RB-TnSeq library were carried out in 2.5L containing 1L of culture volume. Experiments measuring PHB productivity were carried out in 250 mL Ultrayield flasks containing 60 mL of culture volume. Oxygen saturation was verified in all growing conditions using a Fisherbrand Traceable Dissolved Oxygen Pen (part number 15-078-199). Cultures incubated in nitrogen-free medium were first grown for 24 hours in replete DM9 medium containing 1 g/L NH_4_Cl, after which cells were washed three times in DM9 containing 0 g/L NH_4_Cl and resuspended in the same medium. These cultures were then incubated an additional 24 hours to allow for PHB accumulation.

### Culturing in Balanced Growth Conditions to Suppress PHB Production

It was found that frequent subcultures, maintaining an elevated growth rate, was able to suppress PHB accumulation in *C. basilensis* 4G11 (figure S2). This procedure was carried out by culturing cells in fresh medium and incubating for 4-5 generations before passaging the cells. This process was repeated with the culture OD_600_ value being kept perpetually below a value of 0.1, which we found to be the OD at which bulk PHB accumulation begins in our growing conditions (data not shown). This protocol was used to prepare RB-TnSeq library cells for subsequent screening, and to measure PHB productivity of subsequently generated markerless deletion mutant strains.

### Culturing in Unbalanced Growth Conditions to Maximize PHB Production

Cells were inoculated into replete DM9 medium and incubated for 24 hours at 30°C shaking at 250 rpm, yielding stationary phase culture. Cells were then washed three times with DM9 medium in which the NH_4_Cl had been omitted (N-free DM9 medium). Each wash used 45 mL of N-free medium and was done by centrifuging the cells at 15,317 x g at 4°C for 10 minutes. Cells were then resuspended in Nitrogen-free DM9 medium and incubated for an additional 24 hours at 30°C, shaking at 250 rpm.

### Screening *C. basilensis* 4G11 RB-TnSeq Libraries

*C. basilensis* 4G11 RB-TnSeq cells were grown as described above, to prepare replete-grown, low PHB cells in order to screen for putative constitutively producing PHB producing mutants. The cells were then stained using BODIPY 493/503 dye (Invitrogen/ThermoFisher Scientific, part number D3922) to stain intracellular PHB^24^. Stained cells were then analyzed and sorted on a Sony SH800 cell sorter, using the instrument’s 488 nm excitation laser and FITC detection channel and performed in the instrument’s ‘semi-purity’ mode. Gates were constructed around the entire cell population based on forward scattering/side scattering bivariate plots (gate 1). Mutant populations were collected and analyzed from this gate to control for any bias the cell sorter may impart on strain enrichment outside of fluorescence, as done previously^49^. Two other gates were also constructed representing (separately) the 5% most (gate 2) and 5% least (gate 3) fluorescent populations. To sort populations in which PHB accumulation had been suppressed (the ‘low’ PHB population), >5M cytometric events were collected which satisfied gate 1 + gate 2 (most fluorescent, highest PHB). In sorting populations which had been incubated in Nitrogen-free medium (the ‘high PHB’ population), >5M cytometric events were collected which satisfied gate 1 + gate 2 (most fluorescent, highest PHB) and, separately, >5M cytometric events which satisfied gate 1 + gate 3 (least fluorescent, lowest PHB). Cell fractions were sorted into 10 mL of DM9 medium containing 100 ug/mL kanamycin sulfate, transferred to 50 mL total volume of culture and grown until early log phase. 50 mL of this culture was collected and centrifuged at 15,317 x g at 4°C for 10 minutes. Supernatant was decanted and cells were transferred to a 1.5 mL Axygen 1.5 mL microtube (Axygen part number MCT-150-C). The cells were then centrifuged at 21,130 x g for two minutes at room temperature in a benchtop microcentrifuge (model number 5424, Eppendorf). The supernatant was decanted by aspiration and the cells were resuspended in 300 μL of a buffer consisting of 20 mM Tris-HCl (Fisher BioReagent part number BP2475-500), 2 mM sodium EDTA (Millipore Sigma part number E5134-50G), 1.2% Triton X-100 (Millipore Sigma part number X100-5ML), herein referred to as buffer ELB. The cell suspension was then stored at −80°C until DNA extraction was performed.

### Productivity Screening

Screening mutants was done by growing cells as described previously. For replete-grown, PHB-suppressed cultures, samples were collected and analyzed at the end of the 4th, 5th and 6th subcultures/passages. 1 mL samples were collected for PHB analysis, centrifuged at 21,130 x g for two minutes at room temperature in an Eppendorf Centrifuge 5424, and dried overnight at 90°C. Dried pellets were then stored at −20°C. Pellets were processed similar to methods described previously^50^; 500 μL of concentrated H_2_SO_4_ (Millipore Sigma, part number 339741) was added to the pellets and the mixture was heated at 90°C for 30 minutes in a stationary dry bath, vortexing briefly every 10 minutes. The samples were then allowed to cool at room temperature for 30 minutes, after which 500 μL of 5 mM H_2_SO_4_ was added and the mixture was vortexed thoroughly. 1 mL samples were also collected for analysis of supernatant acetate before and after each subculture, to assess the amount of acetate consumed. These samples were also centrifuged at 21,130 x g for two minutes at room temperature in an Eppendorf Centrifuge 5424, filtered through a 0.2 μm PVDF syringe filter (Pall, part number 4406) and stored at −20°C until analysis. Digested samples for PHB analysis as well as filtered supernatants for acetate analysis were analyzed using a Shimadzu Prominence HPLC system equipped with a Biorad HPX 87H chromatography column (Bio-rad Laboratories Inc, part number 1250140) and a diode array detector for UV detection. Both acetate and crotonic acid (the product of the above-mentioned PHB acid digestion) were analyzed at 206 nm. Cell dry weight enumerations were done by passing 45 mL of culture over a pre-dried and pre-weighed 0.22 μm filter followed by two successive washes of 5 mL of milli-Q water. The filters were then dried overnight, allowed to cool to room temperature for one hour and weighed again.

### Promoter Alignment, Candidate Operator Motif Identification and Motif Instance Search

Previously, it was demonstrated that the Acetyl CoA-Synthetase (*acs*) gene is regulated by the CrbR transcription factor/response regulator in *Pseudomonas fluorescens*^40^. The putative LuxR-regulated *acs* homolog in the *C. basilensis* 4G11 genome was identified as the gene with the highest corresponding protein homology to *acs* from *Pseudomonas fluorescens*: RR42_RS13380. 70 bp upstream of the RR42_RS13380 start codon was aligned to genomes from species in the genus *Cupriavidus*. From the set of aligning sequences, one representative sequence was taken from each species (11 species total, shown in SI file ‘RS13380_upstream.txt’ and aligned sequences were inputted to the STREME motif/position weight matrix identification tool^51^. Motifs were identified using default parameters and settings. Among candidate motifs identified, one contained a clear inverted repeat motif sequence to which LuxR family transcription factors in Gammaproteobacteria are known to bind^52^. This motif was then used as input to the FIMO motif instance search tool^53^ in order to search for motif sites within the *C. basilensis* 4G11 genome. The FIMO tool was run using default parameters, yielding a list of individual candidate instances of the identified motif in the *C. basilensis* 4G11 genome.

### Whole Proteome Synteny Mapping and TCS Protein Alignment of *Cupriavidus basilensis* 4G11 and *Cupriavidus necator* H16

Computing proteome synteny between *C. basilensis* 4G11 and *C. necator* H16 was done within KBase^54^, for which the public static narrative can be accessed at https://doi.org/10.25982/123676.7/1880128 (DOI: 10.25982/123676.7/1880128). Briefly, the genomes of both species (RefSeq accession NC_008313 for *C. necator* H16 and RefSeq accession NZ_CP010536 for *C. basilensis* 4G11) were uploaded and subsequently used to create a protein synteny plot (figure S1) using the ‘Compare Two Proteomes’ app. Protein alignments between RR42_RS17055 and RR42_RS17060, and their respective homologs in the *C. necator* H16 genome (see figure 3) were aligned using the ‘Pairwise Align’ function in the Geneious Prime software (version 2022.2.2, https://www.geneious.com). Default values and parameters were used for this analysis.

### Batch Growth, Productivity Modeling and Parameter Fitting

We start by assuming that the data can be fit with a previously validated model^15^ and that O_2_ is always saturated and can thus be neglected in our modeling. We parameter fit each individual subculture of the ΔRR42_RS17055 and ΔRR42_RS17060 mutants from replete-grown subculture screening experiments (figure 2, Table S3, Table S4), assuming each subculture is a batch system.

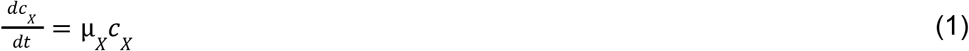

for biomass (*c*_*X*_) and

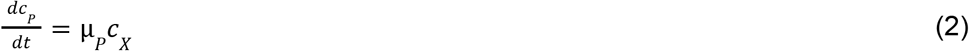

for PHB (*c*_*P*_). In both these equations, μ is the specific growth rate, given as

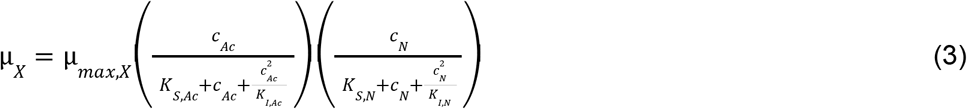

for biomass and

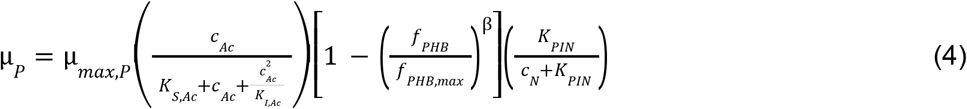

for PHB.

From these equations, the concentrations of acetate and nitrogen (as NH_4_) are

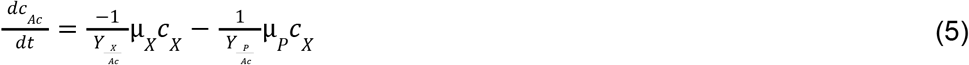

and

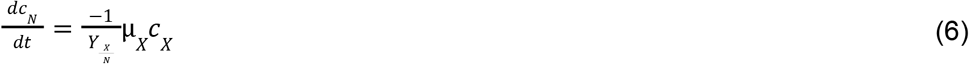

### Parameter Fitting

*K*_*S*,*Ac*_, *K*_*I*,*Ac*_, *K*_*S*,*N*_, *K*_*I*,*N*_, *f*_*PHB*,*max*_, β, and 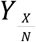 were taken from a previous study^28^.

Initial and final concentrations of acetate, cells, PHB, and nitrogen (as NH_3_) were used to fit remaining parameters 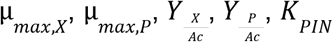, and *K*_*S*,*N*_. Global parameter fitting over each of three batch runs (subcultures 4, 5, and 6) was performed simultaneously. Iterative parameter fitting was run to minimize the following error function using the generalized reduced gradient nonlinear solver function in Microsoft Excel, where c_AC_, c_X_, and c_PHB_ represent the concentration of each of three constituent system components sodium acetate, cell biomass and PHB, respectively.

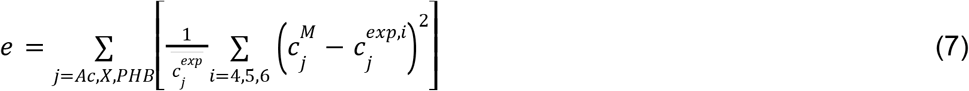

Specific growth rates were constrained to be <0.693 hr^−1^ consistent with that observed previously^28^.

### Continuous Productivity Modeling

All bioreactor models assume well-mixed gas and liquid phases that are exchanged at fixed liquid- and gas-phase dilution rates. In the liquid phase, we consider relevant dissolved constituents which can impact bioproduction^55^; CO_2_, dissolved O_2_, bicarbonate anions (HCO_3_^−^), carbonate anions (CO_3_^2−^), protons (H^+^), hydroxide anions (OH^−^), sodium cations (Na^+^), chloride anions (Cl^−^), acetate anions (H_3_C_2_O_2_^−^), acetic acid (H_3_C_2_O_2_H), ammonia (NH_3_), ammonium cations (NH_4_^+^), microbes (X), and PHB (monomer: C_4_H_6_O_2_).

We consider both a single reactor system and a two-reactor system in which the reactors are connected in series. Equation development for the single reactor is equivalent to that of the first reactor in the two-reactor system. In all following equations, the subscript “*n*” refers to the reactor (*n* has a value either of 1 or 2), and the subscript “*i*” refers to a given species. The reactors are well-mixed and open such that they satisfy mass conservation, given generally for the liquid phase as:

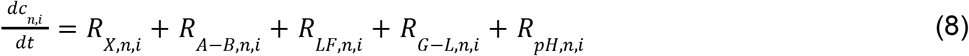

and for the gas phase as:

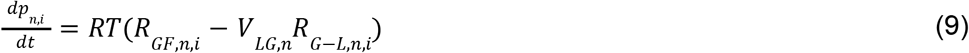

where *c_n,i_* is the concentration; *p_n,i_* is the partial pressure; *R_n,i_* is the net volumetric rate of formation and consumption due to microbial growth (X), acid/base reactions (A–B), liquid or gas flow (LF/GF), gas/liquid mass transfer (G–L), and pH control (pH) for species *i* in reactor *n*; *R* is the gas constant; *T* is the operating temperature; and *V_*LG,n*_* is the ratio of liquid to gas volume in reactor *n* (which we assume to be equal to 2 in all cases).

### Microbial Growth and PHB generation

Microbial growth and PHB generation occur in the well-mixed liquid phase and are responsible for the production of more cells and PHB, and the consumption or production of several chemical species. These reactions are compiled in *R_*X,n,i*_*. Following Mozumder *et al.*^1,2^,

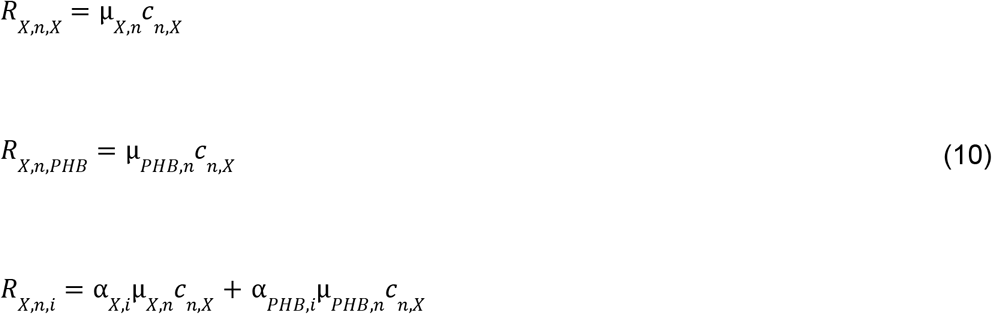

where μ_*X*_ and μ_*PHB*_ are the specific growth rate of cells and the specific accumulation rate of PHB, and α_*X*_ and α_*PHB*_ are stoichiometric coefficients relating biomass growth and PHB accumulation to the consumption or production of other species, as defined below. We set α < 0 if the species is consumed in the reaction. Growth kinetics are described with the inhibition-modified Monod (Andrews-Haldane) model:

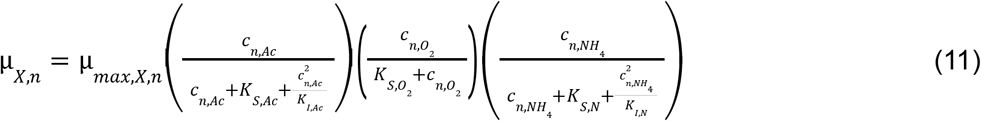

where μ_*max,X*_ is the maximum specific growth rate, and *K*_*S,i*_ and *K*_*I,i*_ are the half-saturation coefficient and inhibition coefficient for species *i*. We modify Mozumder *et al.*^2^ to describe PHB accumulation:

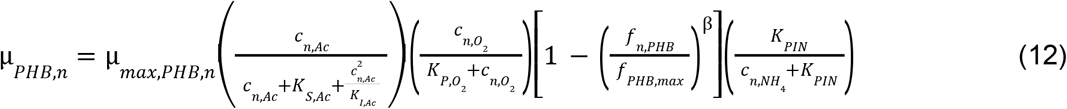

where *f*_*PHB*,(*max*)_ is the (maximum) PHB-to-biomass ratio and *K*_*PIN*_ is the PHB inhibition coefficient for nitrogen.

### Biomass and PHB Yield on Acetate

To determine experimental yields of biomass and PHB on acetate, we first calculated the theoretical yields following the stoichiometric and energetic approach we developed previously^3^. To determine the theoretical yield of biomass on acetate (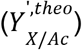), we follow the stoichiometry and energetics of acetate assimilation and oxidation. Acetate is first activated to acetyl-CoA via:

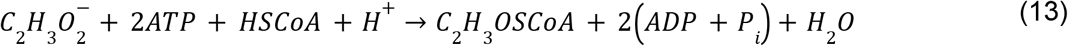

We modify the equation for biomass production from acetyl-CoA developed by Fast and Papoutsakis^56^:

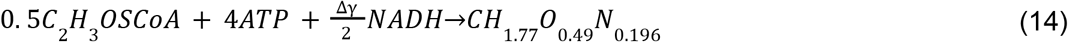

where Δγ is the difference in the degree of reduction between acetyl-CoA (γ = 4) and biomass (γ = 4. 2). We note that this equation, as written, is neither atomically nor charge balanced, so it should be taken to only represent the energy carrier demand of biomass formation. The required energy for acetate activation and acetyl-CoA conversion to biomass is provided by the catabolic reactions of the TCA cycle and by oxidative phosphorylation:

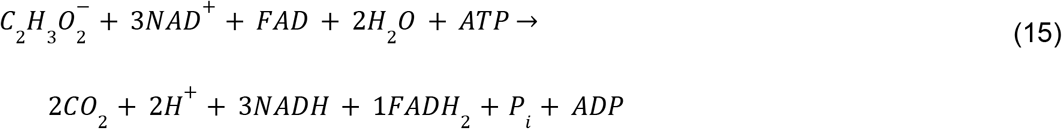

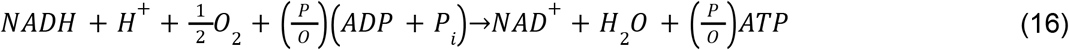

Note that equation (8) includes acetate activation to acetyl-CoA prior to its oxidation in the TCA cycle. We use a *P*/*O* ratio of 2.5 for NADH and 1.5 for FADH_2_ following our previous report^3^. Combining these equations, and including NH_4_^+^ consumption to balance the nitrogen content of biomass, we obtain:

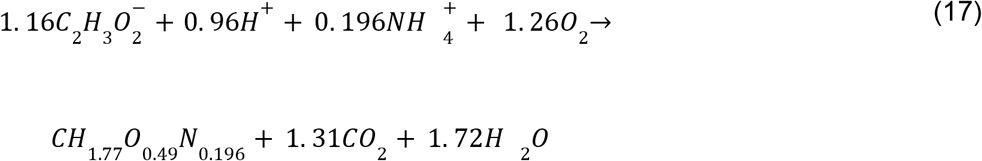

Hence, we use a theoretical molar yield of biomass on acetate of ~0.865 mol mol^−1^ (~0.36 g g^−1^). We use the same approach to calculate the molar yield of PHB on acetate but substitute the biomass-forming equation with the PHB-forming equation, which is given by:

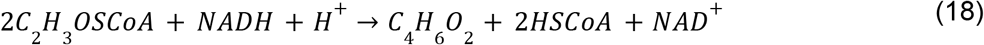

The overall stoichiometry is:

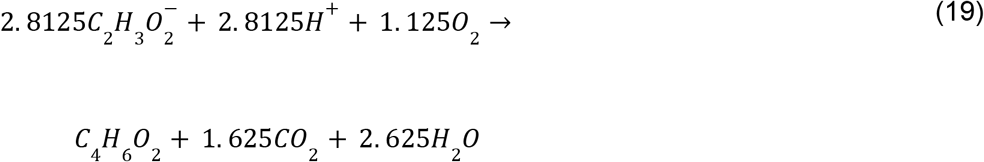

corresponding to a theoretical molar yield of PHB on acetate of ~0.356 mol mol^−1^ (~0.51 g g^−1^). To fit experimental yields of biomass and PHB on acetate, we defined a fractional value, θ_*Y*_, as:

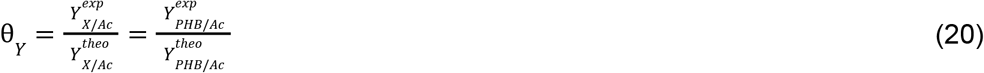

We assume these two ratios are equal on the basis that the metabolic processes producing PHB and cell biomass utilize the same source of carbon (acetate), and the same energy substrate-producing pathways (TCA cycle, oxidative phosphorylation). The production efficiencies of biomass and PHB should thus be nearly equivalent relative to their pathway-constrained maxima.

Hence, in our model, we calculate α_*X*,*n*,*i*_ and α_*PHB*,*n*,*i*_ according to:

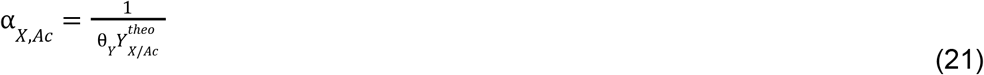

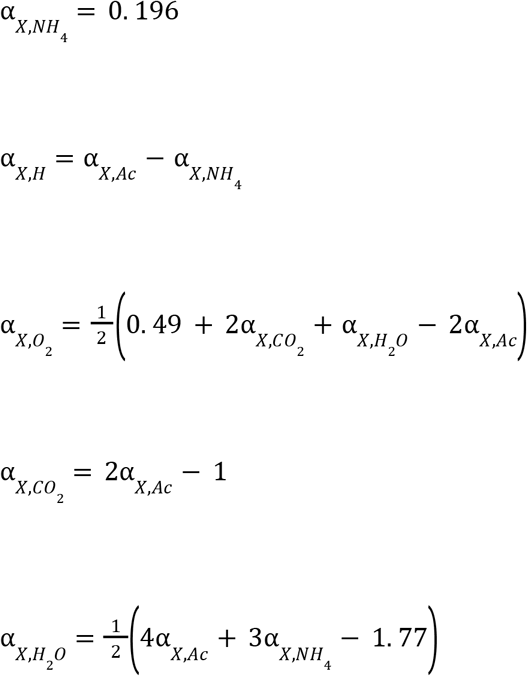

and

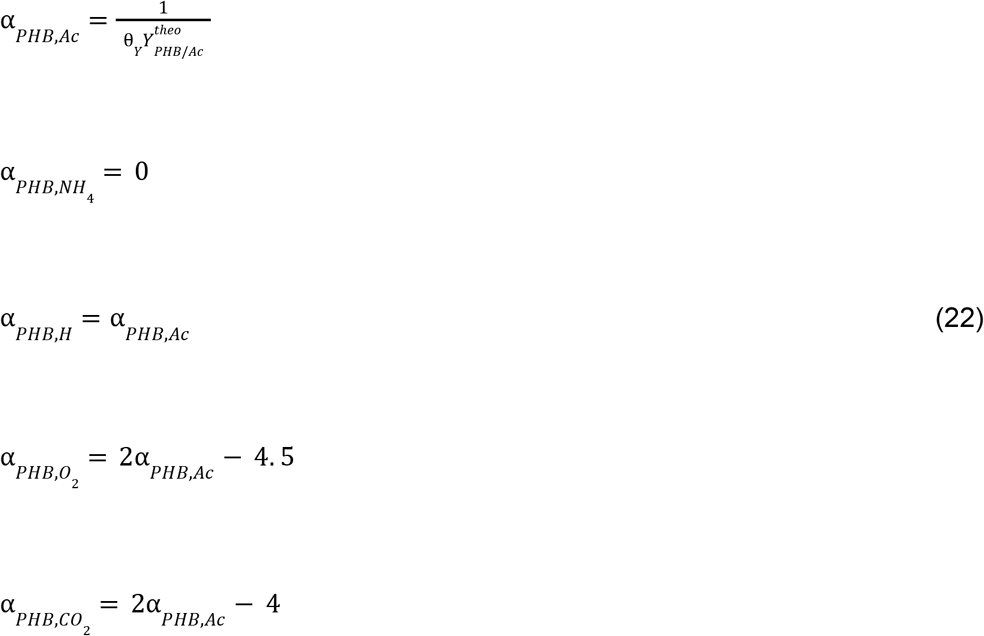

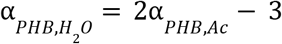

### Growth rate dependence on pH and salinity

We modify our previously developed model to describe the effects of pH and salinity on microbial growth^3^:

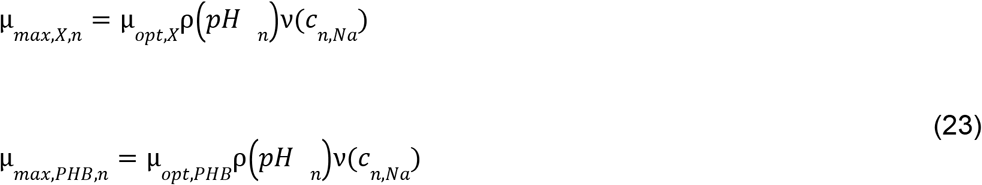

where μ_*opt,X*_ and μ_*opt,PHB*_ are the specific growth and PHB accumulation rates at optimal conditions, and ρ(*pH*) and ν(*c*_*Na*_) are functions describing the impacts of pH and Na^+^ concentration on the growth rate. Following our previous work^3^, we write ρ(*pH*) as:

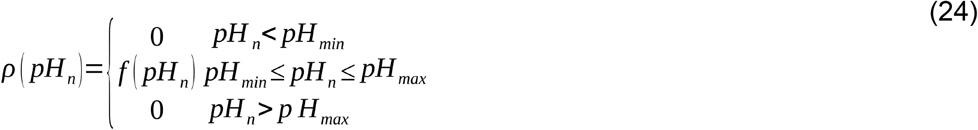

Here, *pH*_*min*/*max*_ is the range of pH over which microbial growth is observed, and the function *f*(*pH_n_*) is:

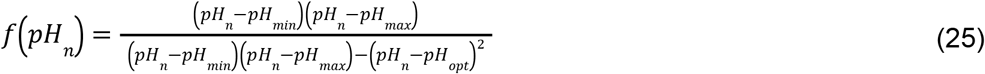

where *pH*_*opt*_ is the optimal pH for growth. Similarly, we define ν(*c_Na_*) as:

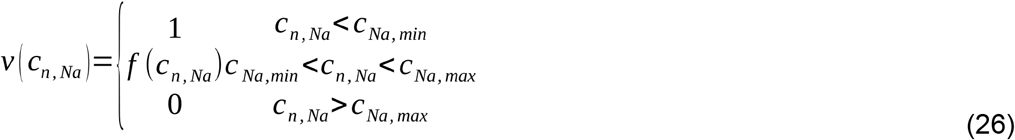

where *c*_*Na,min*/*max*_ is the range of Na^+^ concentration over which growth is impacted, and the function *f*(*c_Na_*) is given by:

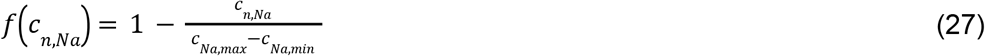

### Acid/base reactions

The acid/base carbon dioxide/bicarbonate/carbonate, acetate/acetic acid, ammonia/ammonium, and water dissociation reactions shown below occur in the liquid phase and are treated as kinetic expressions without assuming equilibrium:

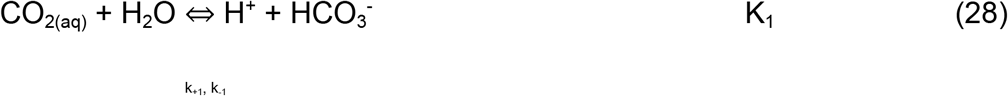

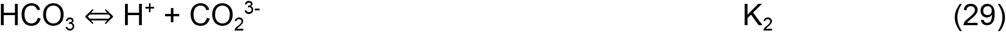

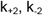

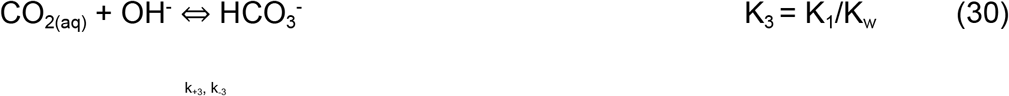

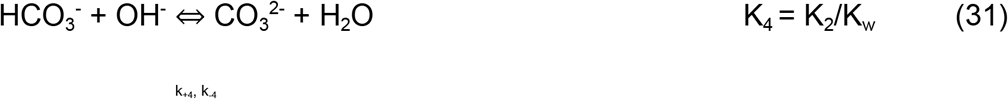

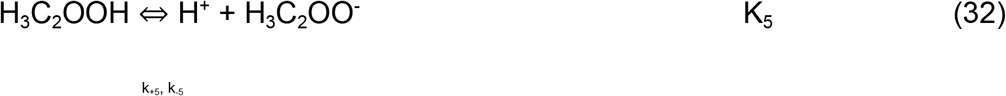

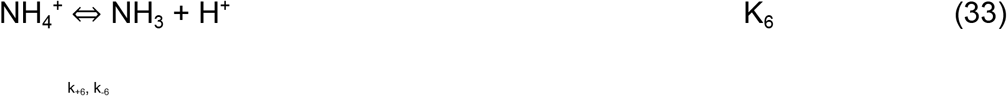

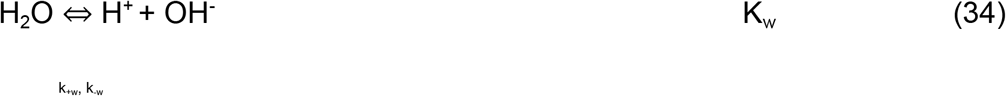

where *k*_+*k*_ and *k*_−*k*_ are the forward and reverse rate constants, respectively, and *K_k_* is the equilibrium constant for the *n*th reaction. For acetic acid and water, we calculate *K_k_* from the van’t Hoff equation using the change of entropy, Δ*S_k_*, and the heat of reaction, Δ*H_k_*, given by:

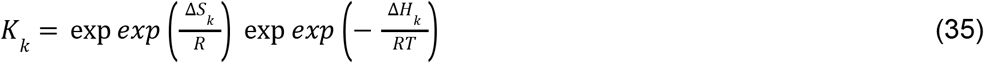

For CO_2_/HCO_3_^−^ and HCO_3_^−^/CO_3_^2−^ equilibria, we calculate *K_k_* using the empirical relationships compiled by W.G. Mook that account for salinity-induced impacts on the equilibrium constant^4^:

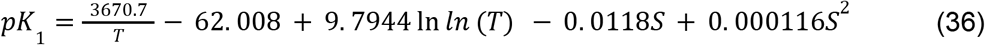

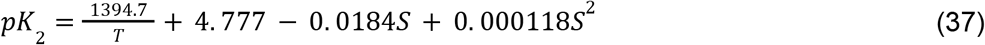

Where *S* is the medium salinity (in units g/kg water).

Source and sink terms resulting from these reactions are compiled in *R*_*A*−*B*,*n*,*i*_, written as:

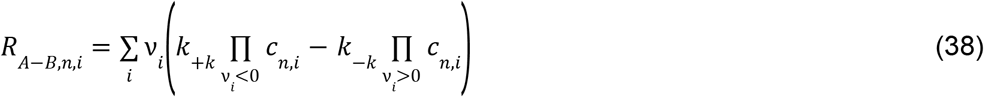

where ν_*i*_ is the stoichiometric coefficient of species *i* for the *k*th reaction and reverse rate constants (*k*_−*k*_) are calculated from:

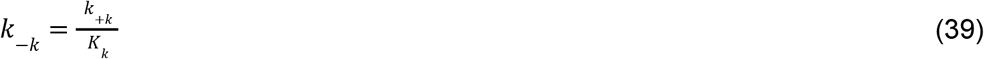

### Liquid and gas flow

Liquid media flows into and out of the well-mixed liquid phase at a constant dilution date (*D*_*liq,n*_). In the first reactor (or lone reactor in the single-reactor system), this results in a feed term written as:

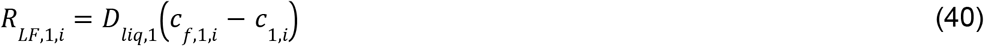

where *c*_*f*,1,*i*_ is the concentration of species *i* in the feed stream to reactor 1. In the two-reactor system, we account for two liquid phase flows. First, the effluent from reactor 1 is fed to reactor 2 directly with no processing. Second, an additional feed stream is optionally applied to supply, for example, additional N-free substrate that can be converted to PHB. On a volumetric basis, then, the total liquid flow rate into the second reactor is the sum of the flow rate out of the first reactor and the additional feed stream flow rate. Hence, the liquid feed term for the second reactor is written as:

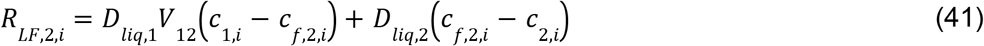

where *V*_12_ is the volumetric ratio of reactor 1 to reactor 2 and *c*_*f*,2,*i*_ is the concentration of species *i* in the optional additional feed stream to reactor 2. Note that when the additional feed stream is not applied, *D*_*liq*,1_*V*_12_ = *D*_*liq*,2_ and *c*_*f*,2,*i*_ = 0 such that the right-hand side of eq. (34) collapses to *D*_*liq*,2_(*c*_1,*i*_ − *c*_2,*i*_) as expected.

For the gas phase(s), we define a feed term according to:

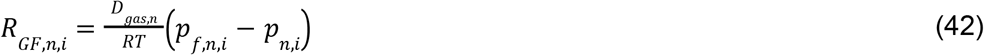

Where *D_gas,n_* is the gas-phase dilution rate in reactor *n*.

### Gas-liquid mass transfer

Gas fed to the reactors result in mass transfer to the liquid phase according to:

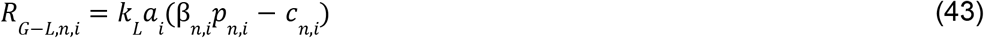

where *k_L_a_i_* us the volumetric mass-transfer coefficient on the liquid side of the gas/liquid interface and β_*n,i*_ is the Bunsen solubility coefficient. We assume 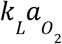 is 300 hr^−1^ following our previous analysis^5^, which represents an intermediate value of the range observed in standard bioreactors^6,7^. To calculate 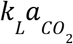, we use:

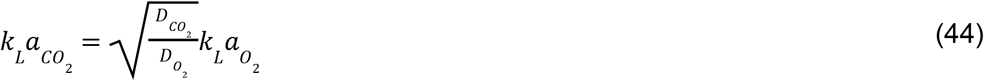

where 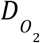 and 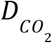 are the diffusivities of O_2_ and CO_2_ following Meraz *et al.*^57^ to account for the differences in the mass transfer coefficient^8^.

We calculate the equilibrium solubility of O_2_ and CO_2_ according to the empirical relationship for the Bunsen solubility coefficient (β):

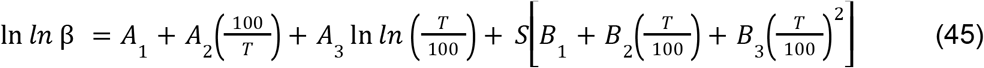

Where *A* and *B* are fitting parameters and *S* is the medium salinity (in units g kg^−1^ water).

### pH control

A feedback control loop is included in the reactor to maintain an optimal pH for microbial growth by adding 1 M sodium hydroxide solution when appropriate. The manipulated flow rate variable (units hr^−1^) is defined as:

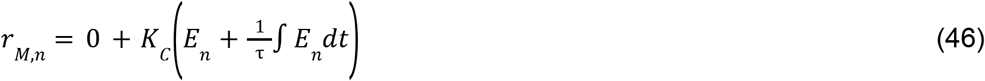

where *K_C_* is the controller gain, *E* is the error, and τ is the controller reset time. The error (*E*) is defined according to:

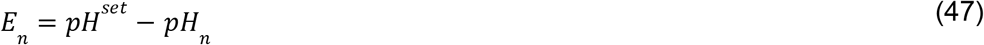

where *pH^set^* is equivalent to *pH_opt_*. The resulting pH control flow is given by:

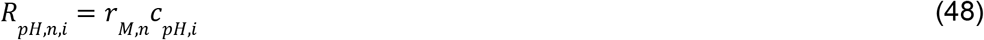

where *c_pH,i_* is 1 M for OH^−^ and Na^+^.

### Model analysis

In the single-reactor system, PHB productivity was calculated simply using:

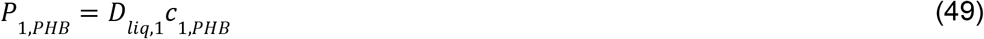

In the two-reactor system, we calculated the overall productivity by normalizing to the combined volume of the reactors, resulting in:

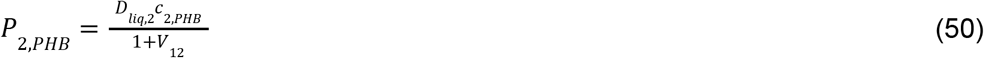

### Model Implementation

All equations were solved using the MUMPS general solver in COMSOL Multiphysics 5.4. Model parameters are listed in Table S2.

### Experimental Reagent and Data Availability

All plasmids used in this study are available through Addgene (https://www.addgene.org/, see table S5 for deposit numbers). All strains used in this study (table S6) are available upon request. Primers used in this study are listed in table S7. The static narrative (DOI: 10.25982/123676.7/1880128) for generating figure S1 is available at https://doi.org/10.25982/123676.7/1880128.

## Supporting information

upstream sequences_RR42_RS13380 homologs

RR42_RS17055 knockout productivity data

RR42_RS17060 knockout productivity data

Table S5_plasmids used in this study

Table S6_primers used in this study

Table S7_strains used in this study

barseq_sample metadata

nFree enrichment raw data

replete enrichment raw data

## Acknowledgements

The authors would like to acknowledge Craig Criddle, Sing Geun Woo, Jorge Meraz, Nils Averesch for invaluable discussion, critique, and feedback during conceptualization and execution of this investigation. We would also like to acknowledge Morgan Price for assistance with fitness determinations as well as gene/protein homology queries and associated discussions. We would also like to acknowledge the Vincent J. Coates Genomics Sequencing Lab at the University of California, Berkeley (QB3 Genomics, UC Berkeley, Berkeley, CA, RRID:SCR_022170) for sequencing mutant amplicon libraries used in this study.

## Author Contributions

**Kyle Sander:** Conceptualization, Data curation, Formal analysis, Investigation, Methodology, Project administration, Validation, Visualization, Writing - original draft, Writing - review & editing. **Anthony J. Abel:** Formal analysis, Methodology, Visualization, Writing - original draft, Writing - review & editing. **Skyler Friedline:** Formal analysis, Investigation, Methodology, Validation, Visualization, Writing - review & editing. **William Sharpless:** Investigation, Methodology, Writing - review & editing. **Jeffrey Skerker:** Conceptualization, Data curation, Project administration, Validation. **Adam Deutschbauer:** Resources, Validation. **Douglas S. Clark:** Methodology, Project administration, Validation, Writing - review & editing, Project administration, Funding acquisition. **Adam P. Arkin:** Conceptualization, Investigation, Formal analysis, Methodology, Project administration, Writing - review & editing, Funding acquisition.

## Citations

1. Kachrimanidou, V. et al. Techno-economic evaluation and life-cycle assessment of poly(3-hydroxybutyrate) production within a biorefinery concept using sunflower-based biodiesel industry by-products. Bioresour. Technol. 326, 124711 (2021).

2. Pavan, F. A. et al. Economic analysis of polyhydroxybutyrate production by Cupriavidus necator using different routes for product recovery. Biochem. Eng. J. 146, 97–104 (2019).

3. Bellini, S., Tommasi, T. & Fino, D. Poly(3-hydroxybutyrate) biosynthesis by Cupriavidus necator: A review on waste substrates utilization for a circular economy approach. Bioresour. Technol. Rep. 17, 100985 (2022).

4. Choi, S. Y. et al. Metabolic engineering for the synthesis of polyesters: A 100-year journey from polyhydroxyalkanoates to non-natural microbial polyesters. Metab. Eng. 58, 47–81 (2020).

5. Kim, B. S. et al. Production of poly(3-hydroxybutyric acid) by fed-batch culture of Alcaligenes eutrophus with glucose concentration control. Biotechnol. Bioeng. 43, 892–898 (1994).

6. Atlić, A. et al. Continuous production of poly(R]-3-hydroxybutyrate) by Cupriavidus necator in a multistage bioreactor cascade. Appl. Microbiol. Biotechnol. 91, 295–304 (2011).

7. Vlaeminck, E., Quataert, K., Uitterhaegen, E., De Winter, K. & Soetaert, W. K. Advanced PHB fermentation strategies with CO2-derived organic acids. J. Biotechnol. 343, 102–109 (2022).

8. Franz, A., Song, H.-S., Ramkrishna, D. & Kienle, A. Experimental and theoretical analysis of poly(β-hydroxybutyrate) formation and consumption in Ralstonia eutropha. Biochem. Eng. J. 55, 49–58 (2011).

9. Lee, S. Y. & Chang, H. N. High cell density cultivation of Escherichia coli W using sucrose as a carbon source. Biotechnol. Lett. 15, 971–974 (1993).

10. Berliner, A. J. et al. Towards a Biomanufactory on Mars. Front. Astron. Space Sci. 8, 120 (2021).

11. Koller, M. & Braunegg, G. Potential and Prospects of Continuous Polyhydroxyalkanoate (PHA) Production. Bioengineering 2, 94 (2015).

12. Vadlja, D., Koller, M., Novak, M., Braunegg, G. & Horvat, P. Footprint area analysis of binary imaged Cupriavidus necator cells to study PHB production at balanced, transient, and limited growth conditions in a cascade process. Appl. Microbiol. Biotechnol. 100, 10065–10080 (2016).

13. Ahn, W. S., Park, S. J. & Lee, S. Y. Production of Poly(3-hydroxybutyrate) by fed-batch culture of recombinant Escherichia coli with a highly concentrated whey solution. Appl. Environ. Microbiol. 66, 3624–3627 (2000).

14. Grousseau, E., Lu, J., Gorret, N., Guillouet, S. E. & Sinskey, A. J. Isopropanol production with engineered Cupriavidus necator as bioproduction platform. Appl. Microbiol. Biotechnol. 98, 4277–4290 (2014).

15. Islam Mozumder, Md. S., Garcia-Gonzalez, L., Wever, H. D. & Volcke, E. I. P. Poly(3-hydroxybutyrate) (PHB) production from CO2: Model development and process optimization. Biochem. Eng. J. 98, 107–116 (2015).

16. Cai, S. et al. A Novel DNA-Binding Protein, PhaR, Plays a Central Role in the Regulation of Polyhydroxyalkanoate Accumulation and Granule Formation in the Haloarchaeon Haloferax mediterranei. Appl. Environ. Microbiol. 81, 373–385 (2015).

17. Juengert, J. R., Patterson, C. & Jendrossek, D. Poly(3-Hydroxybutyrate) (PHB) Polymerase PhaC1 and PHB Depolymerase PhaZa1 of Ralstonia eutropha Are Phosphorylated In Vivo. Appl. Environ. Microbiol. 84, e00604–18 (2018).

18. Juengert, J. R. et al. Absence of ppGpp Leads to Increased Mobilization of Intermediately Accumulated Poly(3-Hydroxybutyrate) in Ralstonia eutropha H16. Appl. Environ. Microbiol. 83, (2017).

19. Klamt, S. & Mahadevan, R. On the feasibility of growth-coupled product synthesis in microbial strains. Metab. Eng. 30, 166–178 (2015).

20. Testa, R. L., Delpino, C., Estrada, V. & Diaz, M. S. Development of in silico strategies to photoautotrophically produce poly-β-hydroxybutyrate (PHB) by cyanobacteria. Algal Res. 62, 102621 (2022).

21. Tang, R., Peng, X., Weng, C. & Han, Y. The Overexpression of Phasin and Regulator Genes Promoting the Synthesis of Polyhydroxybutyrate in Cupriavidus necator H16 under Nonstress Conditions. Appl. Environ. Microbiol. 88, e01458–21 (2022).

22. Price, M. N. et al. Mutant phenotypes for thousands of bacterial genes of unknown function. Nature 557, 503–509 (2018).

23. Ray, J. et al. Complete Genome Sequence of Cupriavidus basilensis 4G11, Isolated from the Oak Ridge Field Research Center Site. Genome Announc. 3, (2015).

24. Kacmar, J., Carlson, R., Balogh, S. J. & Srienc, F. Staining and quantification of poly-3-hydroxybutyrate in Saccharomyces cerevisiae and Cupriavidus necator cell populations using automated flow cytometry. Cytometry A 69A, 27–35 (2006).

25. Price, M., Lo, V. & Shao, W. Fitness Browser.

26. Gu, S. et al. Siderophore-Mediated Interactions Determine the Disease Suppressiveness of Microbial Consortia. mSystems 5, (2020).

27. Shanks, R. M. Q., Caiazza, N. C., Hinsa, S. M., Toutain, C. M. & O’Toole, G. A. Saccharomyces cerevisiae-Based Molecular Tool Kit for Manipulation of Genes from Gram-Negative Bacteria. Appl. Environ. Microbiol. 72, 5027–5036 (2006).

28. Cestellos-Blanco, S. et al. Production of PHB From CO2-Derived Acetate With Minimal Processing Assessed for Space Biomanufacturing. Front. Microbiol. 12, 2126 (2021).

29. Van Wegen, R. J., Ling, Y. & Middelberg, A. P. J. Industrial Production of Polyhydroxyalkanoates Using Escherichia Coll: An Economic Analysis. Chem. Eng. Res. Des. 76, 417–426 (1998).

30. Theodorou, E. C., Theodorou, M. C. & Kyriakidis, D. A. Involvement of the AtoSCDAEB regulon in the high molecular weight poly-(R)-3-hydroxybutyrate biosynthesis in phaCAB+ Escherichia coli. Metab. Eng. 14, 354–365 (2012).

31. Riedel, S. L. et al. Production of poly(3-hydroxybutyrate-co-3-hydroxyhexanoate) by Ralstonia eutropha in high cell density palm oil fermentations. Biotechnol. Bioeng. 109, 74–83 (2012).

32. Ibrahim, M. H. A. & Steinbüchel, A. High-Cell-Density Cyclic Fed-Batch Fermentation of a Poly(3-Hydroxybutyrate)-Accumulating Thermophile, Chelatococcus sp. Strain MW10. Appl. Environ. Microbiol. 76, 7890–7895 (2010).

33. Choi, J. & Lee, S. Y. Factors affecting the economics of polyhydroxyalkanoate production by bacterial fermentation. Appl. Microbiol. Biotechnol. 51, 13–21 (1999).

34. Olavarria, K., Pijman, Y. O., Cabrera, R., van Loosdrecht, M. C. M. & Wahl, S. A. Engineering an acetoacetyl-CoA reductase from Cupriavidus necator toward NADH preference under physiological conditions. Sci. Rep. 12, 3757 (2022).

35. Krauβe, D. et al. Essential Role of the hprK Gene in Ralstonia eutropha H16. J. Mol. Microbiol. Biotechnol. 17, 146–152 (2009).

36. York, G. M., Stubbe, J. & Sinskey, A. J. The Ralstonia eutropha PhaR Protein Couples Synthesis of the PhaP Phasin to the Presence of Polyhydroxybutyrate in Cells and Promotes Polyhydroxybutyrate Production. J. Bacteriol. 184, 59–66 (2002).

37. Pötter, M., Müller, H. & Steinbüchel, A. Influence of homologous phasins (PhaP) on PHA accumulation and regulation of their expression by the transcriptional repressor PhaR in Ralstonia eutropha H16. Microbiology, 151, 825–833 (2005).

38. Karstens, K., Zschiedrich, C. P., Bowien, B., Stülke, J. & Görke, B. Phosphotransferase protein EIIANtr interacts with SpoT, a key enzyme of the stringent response, in Ralstonia eutropha H16. Microbiology, 160, 711–722 (2014).

39. Shi, Y. Serine/Threonine Phosphatases: Mechanism through Structure. Cell 139, 468–484 (2009).

40. Sepulveda, E. & Lupas, A. N. Characterization of the CrbS/R Two-Component System in Pseudomonas fluorescens Reveals a New Set of Genes under Its Control and a DNA Motif Required for CrbR-Mediated Transcriptional Activation. Front. Microbiol. 8, (2017).

41. Garg, R. P., Huang, J., Yindeeyoungyeon, W., Denny, T. P. & Schell, M. A. Multicomponent Transcriptional Regulation at the Complex Promoter of the Exopolysaccharide I Biosynthetic Operon ofRalstonia solanacearum. J. Bacteriol. 182, 6659–6666 (2000).

42. Schell, M. A. Control of Virulence and Pathogenicity Genes of Ralstonia Solanacearum by an Elaborate Sensory Network. Annu. Rev. Phytopathol. 38, 263–292 (2000).

43. Mori, Y. et al. Involvement of ralfuranones in the quorum sensing signalling pathway and virulence of Ralstonia solanacearum strain OE1-1. Mol. Plant Pathol. 19, 454–463 (2018).

44. Lykidis, A. et al. The Complete Multipartite Genome Sequence of Cupriavidus necator JMP134, a Versatile Pollutant Degrader. PLOS ONE 5, e9729 (2010).

45. Barrett, C. F. & Parker, M. A. Coexistence of Burkholderia, Cupriavidus, and Rhizobium sp. Nodule Bacteria on two Mimosa spp. in Costa Rica. Appl. Environ. Microbiol. 72, 1198–1206 (2006).

46. Eng, T. et al. High Throughput Fitness Profiling Reveals Loss Of GacS-GacA Regulation Improves Indigoidine Production In Pseudomonas putida. 2021.02.02.429437 at https://doi.org/10.1101/2021.02.02.429437 (2021).

47. Rajeev, L., Garber, M. E. & Mukhopadhyay, A. Tools to map target genes of bacterial two-component system response regulators. Environ. Microbiol. Rep. 12, 267–276 (2020).

48. Wei, Y.-H. et al. Screening and Evaluation of Polyhydroxybutyrate-Producing Strains from Indigenous Isolate Cupriavidus taiwanensis Strains. Int. J. Mol. Sci. 12, 252 (2011).

49. Coradetti, S. T. et al. Functional genomics of lipid metabolism in the oleaginous yeast Rhodosporidium toruloides. eLife 7, e32110 (2018).

50. Tyo, K. E. J., Jin, Y., Espinoza, F. A. & Stephanopoulos, G. Identification of gene disruptions for increased poly-3-hydroxybutyrate accumulation in Synechocystis PCC 6803. Biotechnol. Prog. 25, 1236–1243 (2009).

51. Bailey, T. L. STREME: accurate and versatile sequence motif discovery. Bioinformatics 37, 2834–2840 (2021).

52. Novichkov, P. S. et al. RegPrecise 3.0 – A resource for genome-scale exploration of transcriptional regulation in bacteria. BMC Genomics 14, 745 (2013).

53. Grant, C. E., Bailey, T. L. & Noble, W. S. FIMO: scanning for occurrences of a given motif. Bioinformatics 27, 1017–1018 (2011).

54. Arkin, A. P. et al. KBase: The United States Department of Energy Systems Biology Knowledgebase. Nat. Biotechnol. 36, 566–569 (2018).

55. Abel, A. J., Adams, J. D. & Clark, D. S. A comparative life cycle analysis of electromicrobial production systems. 2021.07.01.450744 at https://doi.org/10.1101/2021.07.01.450744 (2021).

56. Fast, A. G. & Papoutsakis, E. T. Stoichiometric and energetic analyses of non-photosynthetic CO2-fixation pathways to support synthetic biology strategies for production of fuels and chemicals. Curr. Opin. Chem. Eng. 1, 380–395 (2012).

57. Meraz, J. L., Dubrawski, K. L., El Abbadi, S. H., Choo, K.-H. & Criddle, C. S. Membrane and Fluid Contactors for Safe and Efficient Methane Delivery in Methanotrophic Bioreactors. J. Environ. Eng. 146, 03120006 (2020).

