## Supplemental Information for "Eliminating Genes for a Two Component System Increases PHB Productivity in *Cupriavidus basilensis* 4G11 Under PHB Suppressing, Non-Stress Conditions"

### Title

### Affiliations

[&]Current Affiliation: Department of Bioengineering, University of San Diego, San Diego, CA

[%]Current Affiliation: Zymergen, Emeryville, CA

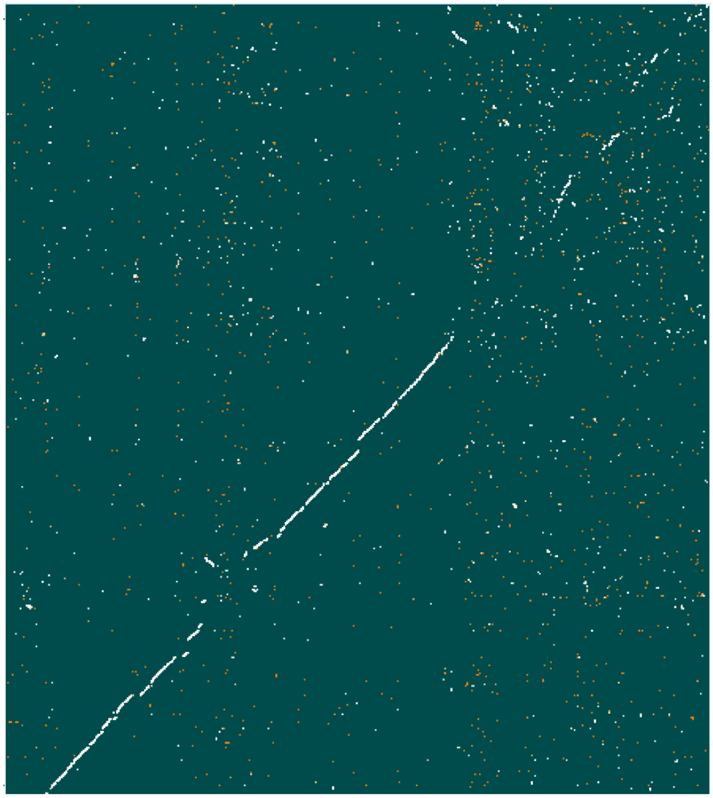

Figure S1: Synteny plot of *C. necator* H16 and *C. basilensis* 4G11 proteomes. The *C. necator* H16 genome is represented along the x-axis and the *C. basilensis* 4G11 genome along the y-axis. This figure was produced using the ‘Compare Two Proteomes’ application which is based on proteins matched as BUS’s (best unambiguous subsets)[1]. White dots indicate protein matches within subsets whose Sub-optimal Best Bidirectional Hit Ratio are >90% and red dots indicate matches whose ratio is <90%.

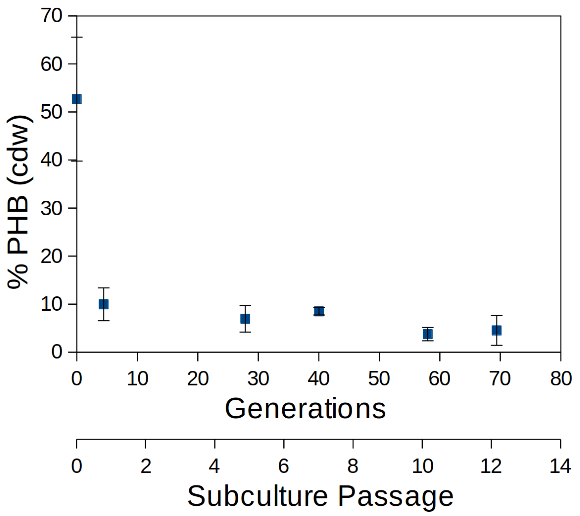

Figure S2: Suppression of PHB accumulation by frequent subculturing. This pre-conditioning strategy was used prior to experiment or analysis throughout this study.

| | | Non-stress Cultivation | | Stress Cultivation | | | | |
|---|---|---|---|---|---|---|---|---|
| | | Enrichment ($>\eta_{95}$) | | Enrichment (FACS) | | | | |
| **Locus Tag** | **Annotation** | **FACS** | **Buoyant Density** | $>\eta_{95}$ | $<\eta_5$ | **Number of Strains** | **Strain Fitness Consistency** | **Average Counts** |
| RR42_RS35075 | AraC family transcriptional regulator | **2.02** | -0.24 | N/A | N/A | 22 | 0.62 | 13.2 |
| RR42_RS02060 | rapZ | **-4.86** | **2.04** | N/A | N/A | 33 | 0.96 | 476.3 |
| RR42_RS28935 | hypothetical protein | **-4.08** | **1.79** | N/A | N/A | 24 | 0.98 | 1382.4 |
| RR42_RS18960 | ABC transporter substrate-binding protein | N/A | N/A | 2.79 | -5.47 | 52 | 0.92 | 518.5 |
| RR42_RS18965 | ABC transporter permease | N/A | N/A | 2.73 | -5.98 | 40 | 0.96 | 611.9 |
| RR42_RS18970 | Toluene ABC transporter ATP-binding protein | N/A | N/A | 2.26 | -6.76 | 40 | 0.98 | 607.3 |
| RR42_RS17060 | Hybrid sensor histidine kinase/response regulator | N/A | N/A | 1.89 | 0.78 | 73 | 0.5 | 108.7 |
| RR42_RS17055 | LuxR family transcriptional regulator | N/A | N/A | 1.61 | 1.18 | 25 | 0.48* | 124.4 |

Table S1: List of genes identified from RB-TnSeq library screening of cells grown in non-stress and nitrogen-limited stress conditions. For non-stress grown libraries, sorted stained cells using FACS (Fluorescence Assisted Cell Sorting) and, separately, by separating cells on buoyant density gradients. For both techniques, fractions representing cells containing the 5% highest PHB accumulation were collected. Nitrogen-limited grown cells were sorted using only FACS, and fractions were collected comprising the 5% highest and, separately, 5% lowest PHB accumulation.

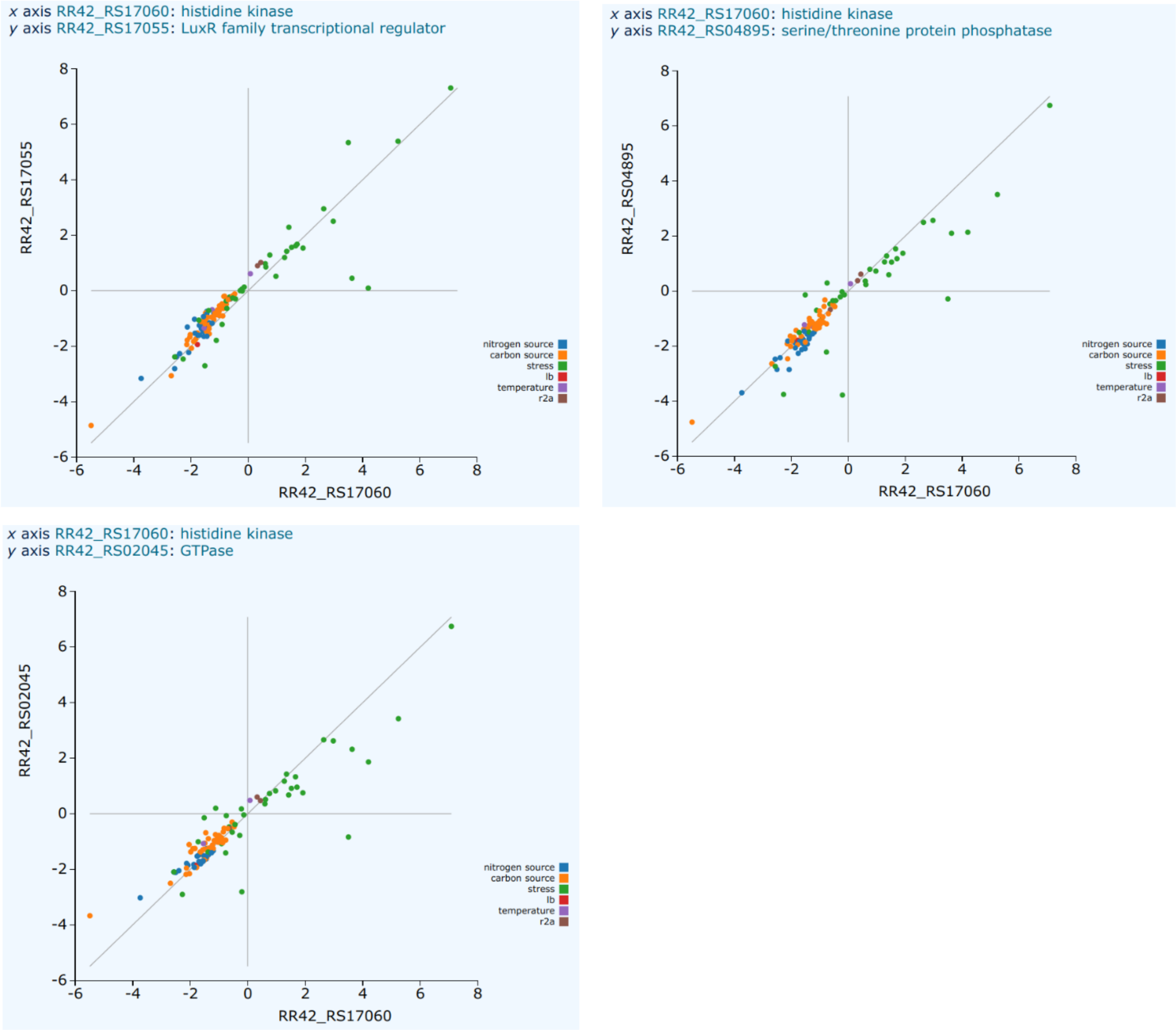


Figure S3: Genes from 106 previous fitness determinations[2] exhibiting similar fitness patterns (using the same *C. basilensis* 4G11 RB-TnSeq library) as the two TCS genes identified in this study. Along the axes of each plot are differential fitness values indicating relative abundance of mutants containing a transposon in the gene indicated along the corresponding axis. Dots indicate these values for each individual experiment. These experiments generally measure differential relative mutant abundance before and after exposure to a stress, antibiotic, or a particular source of carbon or nitrogen in a defined medium setting. Collinearity between two genes indicates similar fitness advantages/defects between knockout mutants of the two genes and can be an indication that the function of two genes are related, part of the same metabolic pathway or signaling cascade. (a) Similar fitness patterns between the two genes comprising the two component system; RR42_RS17055 and RR42_RS17060 (b) Similar fitness patterns exhibited between RR42_RS17060 and a gene annotated as a Serine/Threonine Protein Phosphatase; RR42_RS04895 (c) Similar fitness patterns exhibited between RR42_RS17060 and a gene annotated as a GTPase; RR42_RS02045.

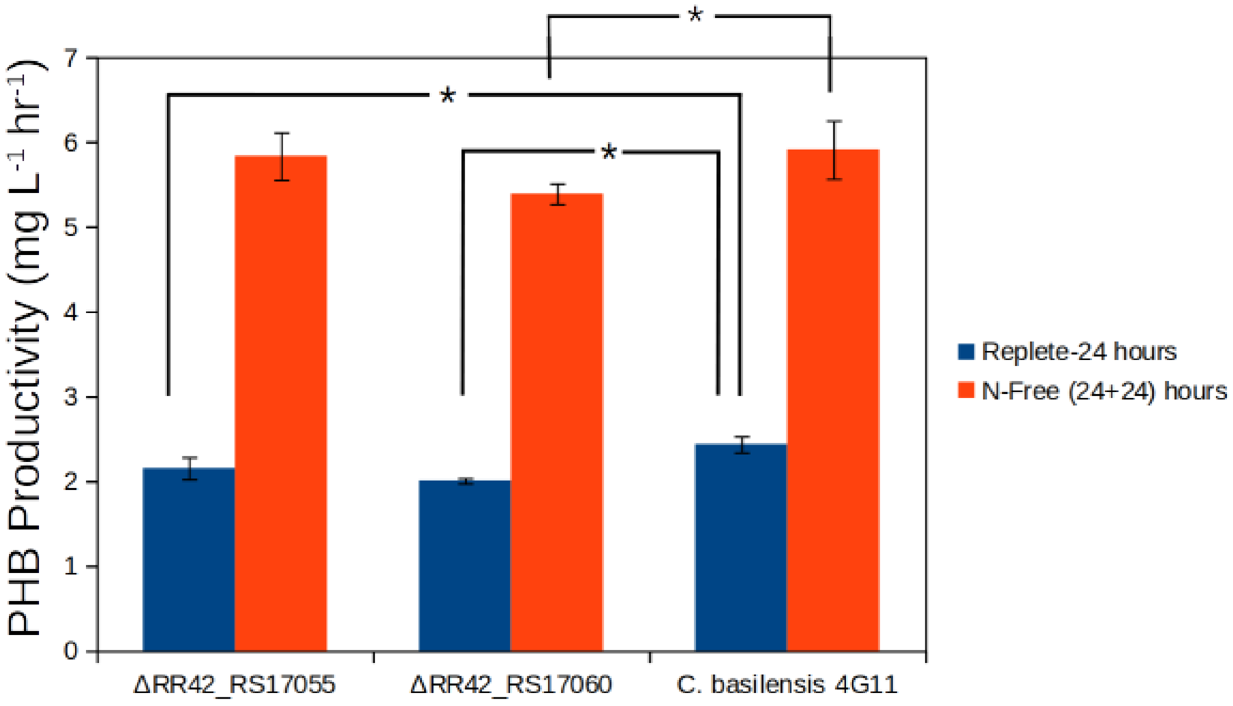


Figure S4: PHB productivity assays of gene deletion mutants of RR42_RS17055 and RR42_RS17055 when grown in defined replete medium, though not subcultured and allowed to grow until stationary phase (blue bars) and after subsequent transfer to nitrogen free defined medium and incubated for an additional 24 hours; both conditions known to induce PHB accumulation in the wildtype organism and other PHB accumulating species. * denotes statistical significance (one tailed t-test, n=3, equal variance assumed, $p<0.05$).

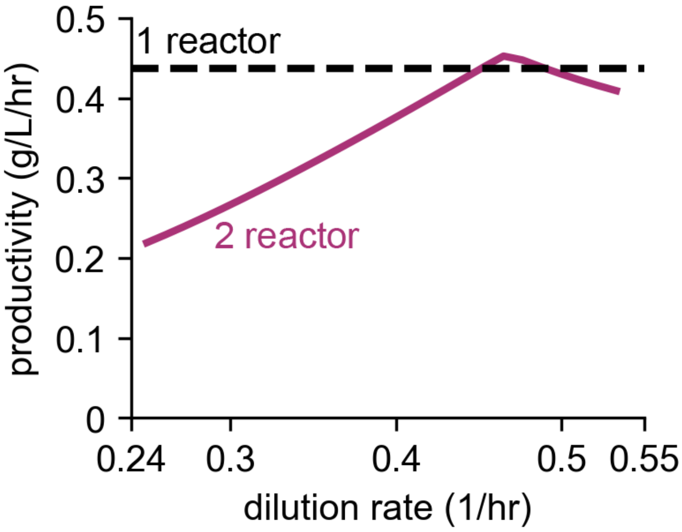


Figure S5: Productivity of *C. basilensis* 4G11 in a single-stage continuous system and a two-stage continuous system.

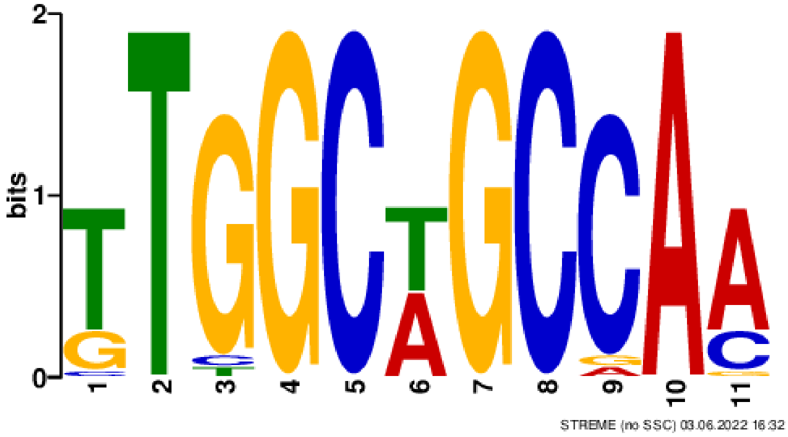


Figure S6: Inverted repeat motif found identified upstream of the gene encoding acetyl-CoA synthetase in *C. basilensis* 4G11 and several other closely related species of *Cupriavidus*. Acetyl CoA synthetase was previously found to be in the regulon of the CrbR regulator; CrbR and the protein encoded by RR42_RS17055 share 34% homology across the entire length of the protein.

Table S2. Model parameters for continuous PHB production.

| Parameter | Value | Description | Units | Source |
|---|---|---|---|---|
| *Reactor & operating conditions* | | | | |
| $T$ | 30 | Temperature | °C | DSMZ |
| $P_{O_2}$ | 0.21 | Partial pressure of $O_2$ | atm | assumed |
| $P_{CO_2}$ | 0.1 | Partial pressure of $CO_2$ | atm | assumed |
| $D_{gas}$ | 100 | Gas phase dilution rate | $hr^{-1}$ | [3] |
| $V_{LG}$ | 2 | Ratio of liquid to gas volume in reactor | -- | assumed |
| $V_{12}$ | 1 | Volumetric ratio of reactor 1 to reactor 2 | -- | assumed |
| $c_{f,1,AAA}$ | 0.167 (10) | Feed concentration of acetic acid to reactor 1 | M (g $L^{-1}$) | assumed |
| | | | M (g $L^{-1}$) | assumed |
| $k_L a_{O_2}$ | 300 | Mass transfer coefficient * interface area available for gas-liquid exchange | $hr^{-1}$ | assumed |
| *Microbial growth* | | | | |
| $\mu_{opt,X}$ | 0.567 | Optimal growth rate of *C. basilensis* 4G11 | $hr^{-1}$ | fitted |
| $Y^{theo}_{X/Ac}$ | 0.36 | Theoretical maximum yield of cell biomass production per gram of acetate | gCDW $g^{-1}$ | calculated |
| $\theta_Y$ | 0.45 | Fraction relating theoretical and experimental biomass and PHB yields, see eqn. 13 | -- | fitted |
| $Y_{X/N}$ | 9.1 | Theoretical maximum yield of cell biomass production per gram of nitrogen | gCDW $g^{-1}$ | [4] |
| $K_{S,Ac}$ | 5 (0.3) | Half-saturation constant - acetate | μM (mg $L^{-1}$) | [5] |
| $K_{I,Ac}$ | 0.83 (49) | Inhibition constant, acetate | M (g $L^{-1}$) | [6] |
| $K_{S,O_2}$ | 2.5 (0.08) | Half-saturation constant - oxygen | μM (mg $L^{-1}$) | [7] |
| $K_{S,N}$ | 9.46 (0.1325) | Half-saturation constant - nitrogen | mM (g $L^{-1}$) | fitted |
| $K_{I,N}$ | 120 (1.691) | Inhibition constant, nitrogen | mM (g $L^{-1}$) | [4] |
| $pH_{opt}$ | 7 | Optimal (setpoint) pH for cell growth/PHB production | -- | DSMZ |
| $pH_{min}$ | 4 | Maximum pH for cell growth/PHB production | -- | [5] |
| $pH_{max}$ | 9.5 | Minimum pH for cell growth/PHB production | -- | [5] |
| $c_{Na,min}$ | 0.2 | Minimum sodium ion concentration at which cell growth is impacted | M | [5] |
| $c_{Na,max}$ | 1.05 | Sodium ion concentration at which cell growth ceases | M | [5] |

| | | | | |
|---|---|---|---|---|
| *PHB production* | | | | |
| $\mu_{opt,PHB}$ | 0.572 | Optimal specific PHB productivity rate | $hr^{-1}$ | fitted |
| $Y^{theo}_{PHB/Ac}$ | 0.516 | Theoretical maximum yield of PHB production per gram of acetate | gPHB $g^{-1}$ | calculated |
| $\theta_Y$ | 0.45 | Fraction relating theoretical and experimental biomass and PHB yields, see eqn. 13 | -- | fitted |
| $K_{P,O_2}$ | 0.57 (.0182) | Inhibition constant of $O_2$ | μM (mg $L^{-1}$) | [4] |
| $K_{PIN}$ | 9.46 (0.1325) – for wildtype<br>120 (1.691) – for ΔRR42_RS17060 mutant | Inhibition constant relating nitrogen concentration to PHB production | mM (g $L^{-1}$) | fitted |
| $f_{PHB,max}$ | 1.78 (6.13) | Maximum ratio of PHB to biomass | mol $mol^{-1}$ (g $g^{-1}$) | [4] |
| $\beta$ | 3.85 | PHB production saturation power coefficient | -- | [8] |
| *Acid-base reactions* | | | | |
| $S_5$ | -92.4 | Specific entropy of acetic acid | J $mol^{-1}$ $K^{-1}$ | [9] |
| $S_w$ | -80.66 | Specific entropy of water | J $mol^{-1}$ $K^{-1}$ | [9] |
| $H_5$ | -0.4 | Specific enthalpy of acetic acid | kJ $mol^{-1}$ | [9] |
| $H_w$ | 55.84 | Specific enthalpy of water | kJ $mol^{-1}$ | [9] |
| $K_6$ | $5.8 \times 10^{-10}$ | Equilibrium constant, see eqn. 33 | mol $L^{-1}$ | [9] |
| $k_{+1}$ | $\exp exp\left(1246.98 - \frac{6\times10^4}{T}\right.$ | Forward kinetic rate constant, see eqn. 28 | $s^{-1}$ | [10] |
| $k_{+2}$ | 59.44 | Forward kinetic rate constant, see eqn. 29 | $s^{-1}$ | [10] |
| $k_{+3}$ | $2.23 \times 10^3$ | Forward kinetic rate constant, see eqn. 30 | L $mol^{-1}$ $s^{-1}$ | [10] |
| $k_{+4}$ | $6.0 \times 10^9$ | Forward kinetic rate constant, see eqn. 31 | L $mol^{-1}$ $s^{-1}$ | [10] |
| $k_{+5}$ | 10 | Forward kinetic rate constant, see eqn. 32 | $s^{-1}$ | assumed |
| $k_{+6}$ | 10 | Forward kinetic rate constant, see eqn. 33 | $s^{-1}$ | assumed |
| $k_{+w}$ | $2.4 \times 10^{-5}$ | Forward kinetic rate constant, see eqn. 34 | L $mol^{-1}$ $s^{-1}$ | [11] |
| *Diffusion coefficients* | | | | |
| $D_{CO_2}$ | $14.68\times10^{-9}\left(\frac{T}{217.206} - 1\right)^{1.}$ [illegible] | Diffusion coefficient of $CO_2$ | $m^2$ $s^{-1}$ | [12] |
| $D_{O_2}$ | $10^\wedge\left[- 8.410 + \frac{773.8}{T} - \left(\right.\right.$ [illegible] | Diffusion coefficient of $O_2$ | $m^2$ $s^{-1}$ | [13] |
| *Bunsen coefficients* | | | | |
| $A_{1,CO_2}$ | -60.2409 | Solubility coefficient for $CO_2$, see eqn. 38 | -- | [14] |
| $A_{2,CO_2}$ | 93.4517 | Solubility coefficient for $CO_2$, see eqn. 38 | -- | [14] |

| | | | | |
|---|---|---|---|---|
| $A_{3,CO_2}$ | 23.3585 | Solubility coefficient for $CO_2$, see eqn. 38 | -- | [14] |
| $B_{1,CO_2}$ | $2.3517 \times 10^{-2}$ | Solubility coefficient for $CO_2$, see eqn. 38 | -- | [14] |
| $B_{2,CO_2}$ | $-2.3656 \times 10^{-2}$ | Solubility coefficient for $CO_2$, see eqn. 38 | -- | [14] |
| $B_{3,CO_2}$ | $4.7036 \times 10^{-3}$ | Solubility coefficient for $CO_2$, see eqn. 38 | -- | [14] |
| $A_{1,O_2}$ | -58.3877 | Solubility coefficient for $O_2$, see eqn. 38 | -- | [15] |
| $A_{2,O_2}$ | 85.8079 | Solubility coefficient for $O_2$, see eqn. 38 | -- | [15] |
| $A_{3,O_2}$ | 23.8439 | Solubility coefficient for $O_2$, see eqn. 38 | -- | [15] |
| $B_{1,O_2}$ | $3.4892 \times 10^{-2}$ | Solubility coefficient for $O_2$, see eqn. 38 | -- | [15] |
| $B_{2,O_2}$ | $1.5568 \times 10^{-2}$ | Solubility coefficient for $O_2$, see eqn. 38 | -- | [15] |
| $B_{3,O_2}$ | $-1.9387 \times 10^{-3}$ | Solubility coefficient for $O_2$, see eqn. 38 | -- | [15] |
| *pH controller* | | | | |
| $K_C$ | 0.1 | Controller gain for pH control | $hr^{-1}$ | -- |
| $\tau$ | 60 | Controller reset time for pH control | s | -- |

### Citations


1. Kellis, M., Patterson, N., Birren, B., Berger, B. & Lander, E. S. Methods in Comparative Genomics: Genome Correspondence, Gene Identification and Regulatory Motif Discovery. *J. Comput. Biol.* **11**, 319–355 (2004).
2. Price, M. N. *et al.* Mutant phenotypes for thousands of bacterial genes of unknown function. *Nature* **557**, 503–509 (2018).
3. Abel, A. J., Adams, J. D. & Clark, D. S. A comparative life cycle analysis of electromicrobial production systems. 2021.07.01.450744 (2021) doi:10.1101/2021.07.01.450744.
4. Islam Mozumder, Md. S., Garcia-Gonzalez, L., Wever, H. D. & Volcke, E. I. P. Poly(3-hydroxybutyrate) (PHB) production from CO2: Model development and process optimization. *Biochem. Eng. J.* **98**, 107–116 (2015).
5. Cestellos-Blanco, S. *et al.* Production of PHB From CO2-Derived Acetate With Minimal Processing Assessed for Space Biomanufacturing. *Front. Microbiol.* **12**, 2126 (2021).
6. Xiao, Y. *et al.* Kinetic Modeling and Isotopic Investigation of Isobutanol Fermentation by Two Engineered Escherichia coli Strains. *Ind. Eng. Chem. Res.* **51**, 15855–15863 (2012).
7. Stolper, D. A., Revsbech, N. P. & Canfield, D. E. Aerobic growth at nanomolar oxygen concentrations. *Proc. Natl. Acad. Sci.* **107**, 18755–18760 (2010).
8. Mozumder, Md. S. I., Goormachtigh, L., Garcia-Gonzalez, L., De Wever, H. & Volcke, E. I. P. Modeling pure culture heterotrophic production of polyhydroxybutyrate (PHB). *Bioresour. Technol.* **155**, 272–280 (2014).
9. Lide, D. R. *CRC Handbook of Chemistry and Physics*. (CRC Press, 2004).
10. Schulz, K. G., Riebesell, U., Rost, B., Thoms, S. & Zeebe, R. E. Determination of the rate constants for the carbon dioxide to bicarbonate inter-conversion in pH-buffered seawater systems. *Mar. Chem.* **100**, 53–65 (2006).

11. Atkins, P. W. *Physical Chemistry*. (Oxford University Press, 1990).
12. Zeebe, R. E. On the molecular diffusion coefficients of dissolved CO2,HCO3-, and CO32- and their dependence on isotopic mass. *Geochim. Cosmochim. Acta* **75**, 2483–2498 (2011).
13. Han, P. & Bartels, D. M. Temperature Dependence of Oxygen Diffusion in H2O and D2O. *J. Phys. Chem.* **100**, 5597–5602 (1996).
14. Directorate-General for Research and Innovation (European Commission), Riebesell, U., Fabry, V. J., Hansson, L. & Gattuso, J.-P. *Guide to best practices for ocean acidification research and data reporting*. (Publications Office of the European Union, 2011).
15. Weiss, R. F. The solubility of nitrogen, oxygen and argon in water and seawater. *Deep Sea Res. Oceanogr. Abstr.* **17**, 721–735 (1970).
